# Cortactin regulates metastatic dormancy of circulating tumor cells by suppressing mTOR-dependent senescence

**DOI:** 10.1101/2025.03.26.645381

**Authors:** Jianyang Hu, Binyu Zhang, Junhao Chen, Guanyin Huang, Hongchao Zhou, Fan Yang, Ke Liu, Shuqian Zheng, Xuefei Liu, Jialing Liu, Hailiang Hu, Lvhua Wang, Jianglin Zhang, Lingyun Dai, Qingfeng Chen, Xinghua Pan, Hongchang Li, Hao Yu, Xin Hong

## Abstract

Metastatic dormancy often refers to the stable cell cycle arrest of disseminated tumor cells (DTCs) at distant sites. However, whether circulating tumor cells (CTCs) in the blood microenvironment can enter a dormant state prior to extravasation and becoming DTCs remains unclear. Using patient-derived melanoma CTC lines and animal explant models (CDX), we identified a previously unrecognized role of the cytoskeletal regulator cortactin (encoded by CTTN) in controlling mTOR/p53-dependent senescence and metastatic dormancy. Cortactin was localized to Rab7-postive endosomes and engaged in late endosomal tethering and homeostasis. The depletion of cortactin resulted in the accumulation of aberrantly enlarged late endosomal aggregates that were positive for Rab7 and mTOR. The mTOR protein complex was accumulated and activated within these abnormal vesicular structures, leading to robust p53 activation through phosphorylation at serine (S) 15 and S33 sites. Consequently, melanoma CTCs underwent G0/G1 cell cycle arrest and entered cellular senescence. This unusual oncogene-induced senescence (OIS) mechanism was characterized by SASP upregulation, β-galactosidase activity, depletion of Ki-67 and Lamin B1, and elevated mitochondrial ROS (mtROS) levels. Notably, a positive feedback loop between p53 and mtROS was essential for maintaining stable senescence in CTCs. In preclinical CDX mouse models, we developed a sequential therapeutic strategy combining cortactin depletion with anti-Bcl-xL senolytic drugs. Such “One-two punch” treatment strategy effectively eliminated viable CTCs and suppressed metastatic tumor growth *in vivo*. Thus, targeting cortactin to induce CTC senescence, followed by senolytic therapy, may represent a promising strategy to block CTC-mediated metastatic progression.

## Introduction

Metastatic dormancy is a critical yet underexplored phase in cancer progression, where disseminated tumor cells (DTCs) remain in a non-proliferative or quiescent state for prolonged periods at distant sites before reactivation to form overt metastases. Dormant DTCs are usually considered as quiescent cells, however, emerging evidence support that DTCs may enter into a senescence-like state, characterized by stable cell cycle withdrawal, mitochondrial dysfunction, β-gal formation, epigenetic reprogramming and secretion of pro-inflammatory cytokines known as senescence-associated secretory phenotype (SASP) ^1–4^. Reactivated DTCs can give rise to metastatic recurrence and eventually lead to cancer mortality years after the primary tumor treatment. However, it remains a major challenge for the detection and characterization of this elusive tumor subpopulation^1,5^. Identifying factors that trigger dormancy and reactivation is of enormous importance for designing interventions to either maintain dormancy indefinitely or eliminate these cells before they progress to full-blown metastatic tumors^6^.

Before arriving distant organs to become DTCs, the pre-metastatic circulating tumor cells (CTCs) in blood must complete extravasation and invasion through the vasculature to reach distant sites. Studies have shown that CTC enumeration and molecular signatures correlate with disease progression and long-term clinical outcome in a variety of cancer types ^7–11^. Given the remarkable shear flow force and other stressors within the blood microenvironmental, most CTCs will undergo apoptosis and a tiny fraction of them remain viable and proceed to become DTCs. It is widely believed that metastatic dormancy occurs in DTCs that originated from the few surviving CTCs. However, it remains unclear whether CTCs can enter dormancy before extravasation.

Cellular senescence is originally defined as an irreversible state of withdrawal from the cell cycle, induced by intrinsic stressors such as DNA replicative stress or damage, oncogene activation, oxidative stress, and telomere dysfunction ^12–15^ or environmental stimuli including chemotherapeutic agents, UV light, ionizing radiation and others ^16,17^. Oncogene-induced senescence (OIS), a type of premature senescence, is considered a fail-safe mechanism used to prevent the malignant transformation of cells with uncontrolled proliferation or increased genomic instability ^18^. Nevertheless, OIS evasion has become a hallmark trait of malignant progression in aggressive cancers like melanoma. More than 80% melanocytic nevi harboring BRAF^V600E^ mutation exhibited OIS, characterized by high level of p16ARF proteins and the activation of p53/p21 pathways, serving as a barrier for oncogenic transformation ^19–21^. CDKN2A (p16INK4a, p14ARF) and CDKN2B were frequently lost in melanoma, which is responsible for the melanic nevus to bypass OIS and transform into malignant melanoma ^22^.

Cortactin, encoded by CTTN, is a class 2 nucleation-promoting factor (NPF) which can synergize with class 1 NPFs (such as WAVE, WASP) to stimulate efficient Arp2/3- mediated formation of actin branches^23^. Initially identified as a substrate for Src tyrosine kinases, cortactin has been implicated in various cellular processes, including cell adhesion, migration, invasion, and endocytosis ^23–25^. Genomic amplification of the CTTN gene is common in head, neck, esophageal squamous cell carcinomas and cutaneous melanoma ^26,27^. It is reported that cortactin promotes melanoma cell migration and invasion by enhancing actin polymerization, invadopodia formation, and matrix degradation ^28,29^. Moreover, cortactin interacts with other signaling molecules implicated in melanoma progression, such as integrins, matrix metalloproteinases (MMPs), and growth factor receptors, further augmenting its pro-metastatic effects ^30–32^. As a cytoskeleton regulator, whether and how cortactin is involved in cellular senescence and metastatic dormancy remains a mystery.

This study identified cortactin as one of the highly elevated genes in melanoma CTCs, and its high expression was significantly correlated with poor clinical outcome. Cortactin played a critical role in the senescence induction of melanoma CTCs. Cortactin was co-localized with Rab7 in the late endosomes. Cortactin deletion (CTTN KD) resulted in the formation of large Rab7^+^ mTOR^+^ late endosomal aggregates. The accumulation and overactivation of mTOR activity within these abnormally endosomal vesicular structures led to p53 phosphorylation and stabilization, and consequently senescence induction, marked by G0/G1 cell cycle arrest, SASP upregulation, and defective mitochondrial fission dynamics. The dysregulation of mitochondrial oxidative function resulted in dramatic mtROS accumulation, which further strengthened the activation of p53, forming a signal amplification loop through the“p53 ---| CDK1→p-DRP1 ---| mtROS→p53” axis. Finally, a “One-two punch” sequential treatment strategy using CTTN KD followed by anti-Bcl-xL senolytics was developed to suppress CTC-mediated metastatic progression in preclinical CDX models.

## Results

### Cortactin depletion induces senescence of *ex vivo*-cultured circulating melanoma cells

We integrated smart-seq2 of CTC from 8 public datasets, including skin cutaneous melanoma (SKCM), breast cancer (BRCA), castration-resistant prostate cancer (CRPC) and castrate-sensitive prostate cancer (CSPC) ^33–39^. After quality control and batch correction, 291 CTCs were subjected for unsupervised clustering analyses and 4 distinct CTC subpopulations were identified, including CTTN^high^ CTC (highly expressing CTTN and TET2), HBB^high^ CTC (highly expressing HBB), KRT18^high^ CTC (highly expressing KRT8 and KRT18), and PF4^high^ CTC (highly expressing PF4 and PPBP) (Fig. 1a, Extended Data Fig. 1a,b). The proportions of these 4 CTC subpopulations in different cancer types were categorized and quantified (Fig. 1b). Notably, the CTTN^high^ CTC population was primarily enriched in SKCM, suggesting that this subpopulation might be a distinctive feature representative of SKCM. Higher expression of CTTN in clinical patient samples was associated with poorer overall survival, and distant metastasis-free survival of SKCM patients (Fig. 1c and Extended Data Fig. 1c,d). The elevation of CTTN in melanoma CTCs and its clinical prognostic value promoted us to test if CTTN plays a functionally important role in CTC survival and metastatic progression.

**Fig. 1.**
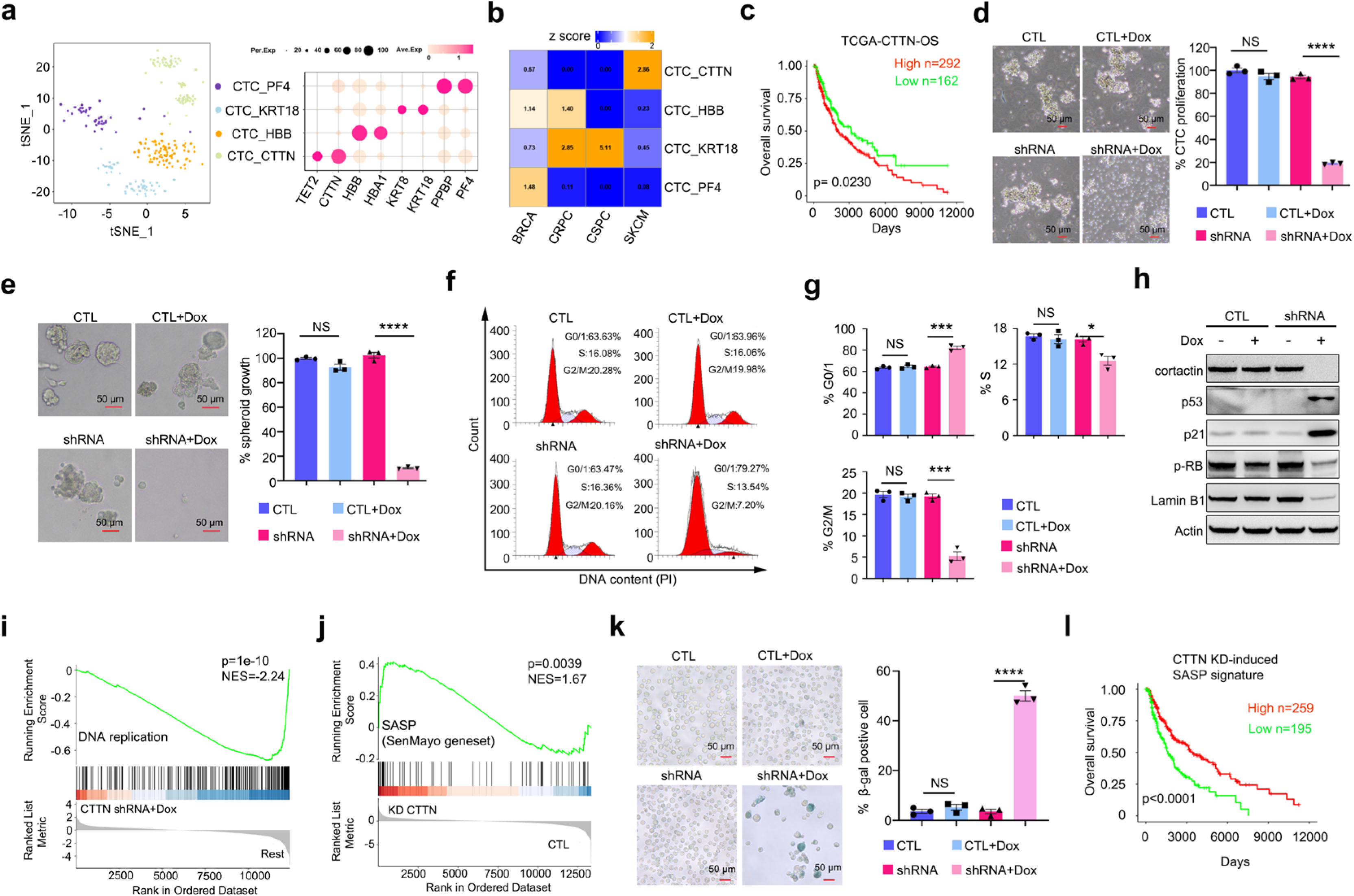
Cortactin depletion induces cellular senescence in *ex vivo*-cultured circulating melanoma cell. **a**, Left panel, t-SNE map showing 4 CTC subpopulations. Right panel, the expression levels of the selected markers in different primary CTC subpopulations. Dot size indicates the fraction of expression cell, and the colors represent normalized gene expression levels. **b,** Quantification of cancer type prevalence of primary CTCs estimated by Ro/e score. **c,** Overall survival of patients with SKCM, analyzed by using the TCGA-SKCM data set. Patients from stage I to IV were all included. Stratification by high and low CTTN mRNA expression. n= 292 and 162 patients with high and low CTTN expression, respectively. **d**, Quantification of cell proliferation of Mel-167 CTC cells cultured in suspension with or without doxycycline-mediated inducible CTTN KD. Pictures showed the representative morphology of different cell lines. n=3 independent experiments. Scale bars, 50 μm. **e**, Quantification of cell proliferation of the different Mel-167 CTC cell lines cultured in a 3D model with or without doxycycline-mediated inducible CTTN KD. Pictures showed the representative morphology of different cell lines. n=3 independent experiments. Scale bars, 50 μm. **f**, Cell cycle analysis of Mel-167 treated with or without doxycycline for 3 days. **g**, Quantification of G0/G1, S, and G2/M phase of cells in (**d)**. n=3 independent experiments. **h**, Immunoblotting detection protein level of cortactin and senescence marker, p53, p21, phosphorylated-RB (p-RB) and Lamin B1. **i,j**, Gene set enrichment analysis showed that the downregulation of DNA replication (**i**), and the upregulation of Senescence-Associated Secretory Phenotype (SenMayo gene set) (**j**) in CTTN KD group compared with control group based on bulk RNA-seq. Doxycycline treatment lasted for 3 days. **k**, Representative picture showed the SA-β-gal staining of Mel-167 cells with or without CTTN KD for 6 days. Quantification of the percentage of SA-β-gal positive cell. n=3 independent experiments. **l**, Kaplan-Meier estimation of survival time of TP53-WT patients in The Cancer Genome Atlas (TCGA)-SKCM dataset based on the expression level of CTTN KD- induced SASP signature. P value was calculated by log rank test. Dox, doxycycline; CTL, doxycycline inducible control shRNA; shRNA, doxycycline inducible CTTN-targeting shRNA. Data are represented as mean ± SEM in (**d**), (**e**), (**g**) and (**k**). The statistical significance was calculated by two-sided Student’s t test in (**d**), (**e**), (**g**), (**k**).\**P*<0.05; \*\*\**P*<0.001; **** *P*<0.0001; NS, not significant. *P*-values were calculated by log-rank test in (**c**)and (**k**).

Using melanoma patient-derived CTC lines ^39^, we constructed doxycycline inducible shRNA system to deplete cortactin expression (CTTN KD) in Mel-167 CTC line and metastatic melanoma A375 cell line (Extended Data Fig. 1e). The cell proliferation in suspension (Mel-167) or adherent (A375) and 3D Matrigel (both cell lines) culture models were significantly reduced upon CTTN KD in both cell lines (Fig. 1d,e and Extended Data Fig. 1f,g). Flow cytometry cell cycle analysis showed that the fractions of G0/G1 were significantly increased, while the S and G2/M phases of Mel-167 CTCs were remarkably decreased (Fig. 1f,g). Similar results were observed in A375 cells (Extended Data Fig. 1h). Thus, cortactin positively regulated CTC proliferation and cell cycle progression.

We reasoned that the G0/G1 cell cycle arrest might be a signature of CTC senescence induction upon cortactin depletion. To test this hypothesis, multiple markers in combination were examined to confirm the senescence phenotype, including p53, p21, Lamin B1, phosphorylated Retinoblastoma (p-RB), Ki-67, and beta-gal staining ^40,41^. Strikingly, the depletion of cortactin in Mel-167 and A375 cells led to a significant elevation of p53 and p21, while Lamin B1 and p-RB (inactivated form of RB) were remarkably reduced (Fig. 1h and Extended Data Fig. 1i). RNA-seq analysis was performed to analyze the transcriptomic differences in Mel-167 CTCs with or without CTTN KD (Extended Data Fig. 1j). GSEA analysis showed that the DNA replication pathway was downregulated in CTTN-KD CTCs, in line with the experimental cell cycle analysis (Fig. 1i). The canonical proliferation marker Ki-67 mRNA level was decreased upon CTTN KD (Extended Data Fig. 1k), which was consistent with the overall suppression of E2F targets by bulk RNA-seq analysis in both cell lines (Extended Data Fig. 1l and Extended Data Fig. 4a-c). High expression of CTTN was positively correlated with E2F targets expression in melanoma patient samples (Extended Data Fig. 1m). Senescence-associated secretory phenotype (SASP) is considered a molecular hallmark of senescent cells. GSEA analysis showed that the reported SenMayo SASP gene set ^42^ were significantly enriched in Mel-167 CTC with CTTN-KD compared to the control group (Fig. 1j, Extended Data Fig. 1n). It was reported that the degree of canonical senescence biomarker overexpression were positively correlated with the duration of cell cycle withdrawal ^43^. We compared the SASP gene expression by knocking down CTTN in Mel-167 for 3 days and 4 days, and found that SASP-related gene expression were further elevated in CTTN-KD CTCs for 4 days as compared to CTTN-KD for 3 days (Extended Data Fig. 5e), suggesting these CTCs may gradually enter a deeper senescence state when cortactin was depleted over time. Consistently, another classical senescence marker, the β-galactosidase (β-gal), also showed positivity in CTTN-KD Mel-167 and A375 cells (Fig. 1k and Extended Data Fig. 1o,p). High SASP signature was positively correlated with a better overall survival in the TCGA melanoma patient dataset (Fig. 1l), as well as in another dataset of melanoma patients receiving immune check point blockade therapy (Extended Data Fig. 1q,r).

Collectively, these data indicated that cortactin was critically involved in suppressing senescence induction in both pre-metastatic Mel-167 CTCs and metastatic A375 melanoma cells.

### Heterogenous expression of CTTN are inversely correlated with the senescence state in early cultures of single CTCs

To test if CTTN expression heterogeneity marks CTC proliferation *versus* (*VS*) senescence states, we performed single-cell transcriptomic analysis on early cultures of single Mel-167 CTCs. We scored the expression of G2M- and S-phase genes in CTTN^high^(Top 25%, n=147) and CTTN^low^(Down 25%, n=147) CTCs, followed by grouping them into the cycling CTCs and non-cycling CTCs (Extended Data Fig. 2a) based on the method published by Jakab et al ^44^. Our analysis indicated that the cycling CTCs were predominantly CTTN^high^ subset, taking up 60.2% compared to 39.8% as CTTN^low^ group, while non-cycling CTCs were mainly composed of CTTN^low^ group (67.6% CTTN^low^ *VS* 32.4% CTTN^high^, p=7.6e-06, Extended Data Fig. 2b). Furthermore, single-sample gene set enrichment analysis (ssGSEA) demonstrated that CTCs in CTTN^low^ group exhibited significant elevation of classical senescence-associated signatures, including cellular senescence, the p53 pathway, SASP signatures, mitochondrial electron transportation chain (ETC) pathway, oxidative phosphorylation, ROS signaling, and other signatures indicative of cell cycle arrest compared to CTTN^high^ CTC (Extended Data Fig. 2c,d). Thus, CTTN expression was heterogenous in early cultures of polyclonal CTCs that appeared to be inversely correlated with the expression of senescence signatures. CTTN^low^ CTCs may exist in a pre-senescent state that could be actively primed to enter senescence upon appropriate stimulus.

### Cortactin promotes tumor growth, CTC generation and metastatic colonization

To test if CTTN KD affect the tumorigenic and metastatic potential of CTCs, we established CTC-derived animal explant (CDX) models and found that the inducible CTTN KD at 19 days post subcutaneous injection could efficiently suppress tumor growth, and the tumor-suppressive effect was further enhanced when the knockdown was induced at day 7 using doxycycline (Fig. 2a,b and Extended Data Fig. 3a). CTC- mediated metastasis potency was assessed using the tail-vein injection of Mel-167 CTCs to immunocompromised NCG mice ^39^. Remarkably, inducible CTTN KD post day 1 and day 7 of CTC injection effectively suppressed metastasis formation (Fig. 2c,d,f; Extended Data Fig. 3b-d), which was not due to the non-specific effect of doxycycline addition (Extended Data Fig. 3e-g). Interestingly, the number of circulating secondary CTCs within the mice’s blood was also diminished in the group of CTTN-KD CTCs compared with the control (Fig. 2e). We further assessed if CTTN-KD Mel-167 tumors undergo senescence *in vivo*, we performed immunohistochemistry (IHC) and qPCR to determine the expression of cellular senescence markers and SASP signatures. Our IHC data showed that the p53 and p21 proteins were significantly upregulated, while p-RB protein and Lamin B1 mRNA were downregulated in CDX tumors with CTTN KD (Fig. 2g-i). SASP genes, such as CXCL8, ICAM1, GDF15, FAS, PLAT were also significantly upregulated in CTTN-KD tumors (Fig. 2j).

**Fig. 2.**
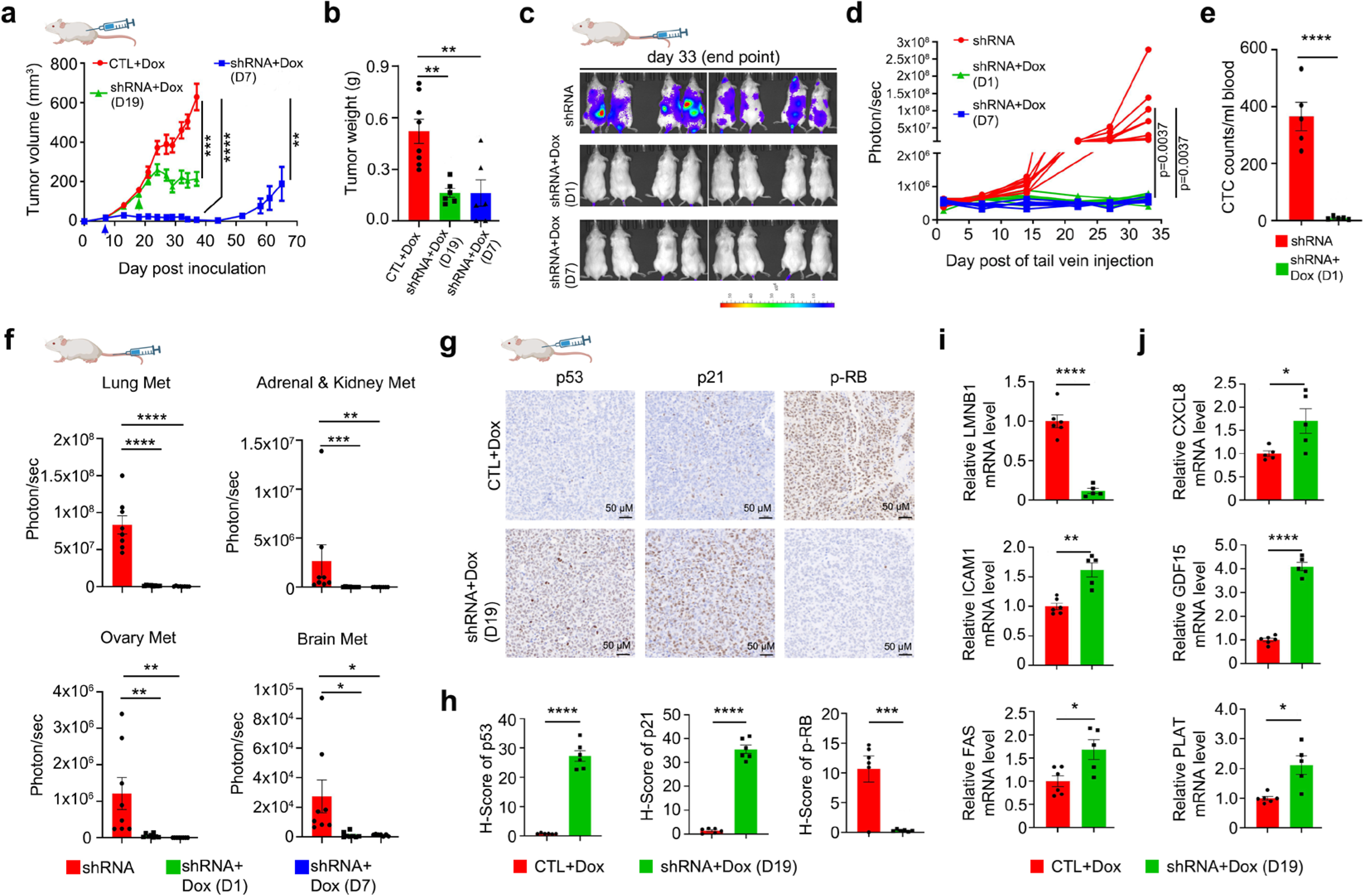
CTTN regulates melanoma tumorigenesis, CTC generation, and metastatic colonization. **a,** Tumor growth curve of Mel-167 CTC subcutaneously injected to NCG mice. n=8 for CTL+Dox group, n=6 for other two groups. The arrow heads indicate the start of doxycycline administration. **b**, Tumor weight at the experiment endpoint. n=8 for CTL+Dox group, n=6 for other two groups. For CTTN shRNA+Dox at day 7, two tumors disappeared at the endpoint, and their weight was recorded as zero. **c**, Tumor metastasis experiment established by injecting Mel-167 CTC expressing luciferase-GFP to NCG mice through the tail vein. Picture shows the IVIS images captured at the experiment endpoint. n=8 mice. **d**, Growth curve of metastases indicated by luciferase signal of whole mouse detected by IVIS system. n=8 mice. **e**, GFP-positive CTC number. Cells were isolated from 1 ml blood of the metastasis mouse model by the Celutriator TX1 microfluidic device. n=5 mice **f,** Bar graphs showed the quantification of ex vivo luciferase signal of metastases from the lung, adrenal and kidney, ovary, and brain collected from (**c**). n=8 organs. **g**, Representative IHC pictures of p53, p21, and phosphorylated-RB ser807/811 in Mel-167 s.c. xenograft model. Scale bar, 50 μm. **h**, Quantification of p53, p21 and phosphorylated-RB expression of (**g**). n=6 tumors. **i**, LMNB1 (Lamin B1) mRNA level in Mel-167 xenograft model detected by qPCR. n=6 tumors. **j**, SASP genes, CXCL8, ICAM1, GDF15, FAS, and PLAT mRNA level in Mel-167 xenograft model detected by qPCR. n=6 tumors. Data are represented as mean ± SEM in (**a**), (**b**), (**e)**, (**f**) and (**h-j**). The statistical significance was calculated by two-sided Student’s t test in (**a**), (**b**), (**d**-**f**) and (**h-j**). \**P*<0.05; \*\**P*<0.01; \*\*\**P*<0.001; **** *P*<0.0001.

Collectively, these data suggested that CTTN KD triggered tumor senescence *in vivo*, which consequently led to marked suppression of melanoma growth, CTC generation, and metastatic colonization.

### Cortactin depletion-induced senescence is p53-dependent

To determine which molecular features are remodeled upon CTTN KD, simultaneous RNA-sequencing (Extended Data Fig. 1j) and proteomic profiling (Fig. 3a) were applied to Mel-167 CTCs with or without CTTN KD. “p53 pathway” and “Oxidative phosphorylation” pathway signatures were positively correlated with CTTN KD while “Mitotic spindle”, “G2M checkpoint” and “E2F targets” signatures were negatively correlated with CTTN KD (Extended Data Fig. 4a,b) in both transcriptomic and proteomic levels. A similar result from RNA-seq was found in A375 cells with CTTN KD (Extended Data Fig. 4c). Classical p53-activated target genes ^45^ such as CDKN1A, MDM2, BAX, GADD45A, BTG2 and PLK3 were significantly upregulated (Extended Data Fig. 4d,e), consistent with the GSEA analysis showing that the p53 pathway signature is highly activated in CTTN-KD CTCs (Fig. 3b,c).

**Fig. 3.**
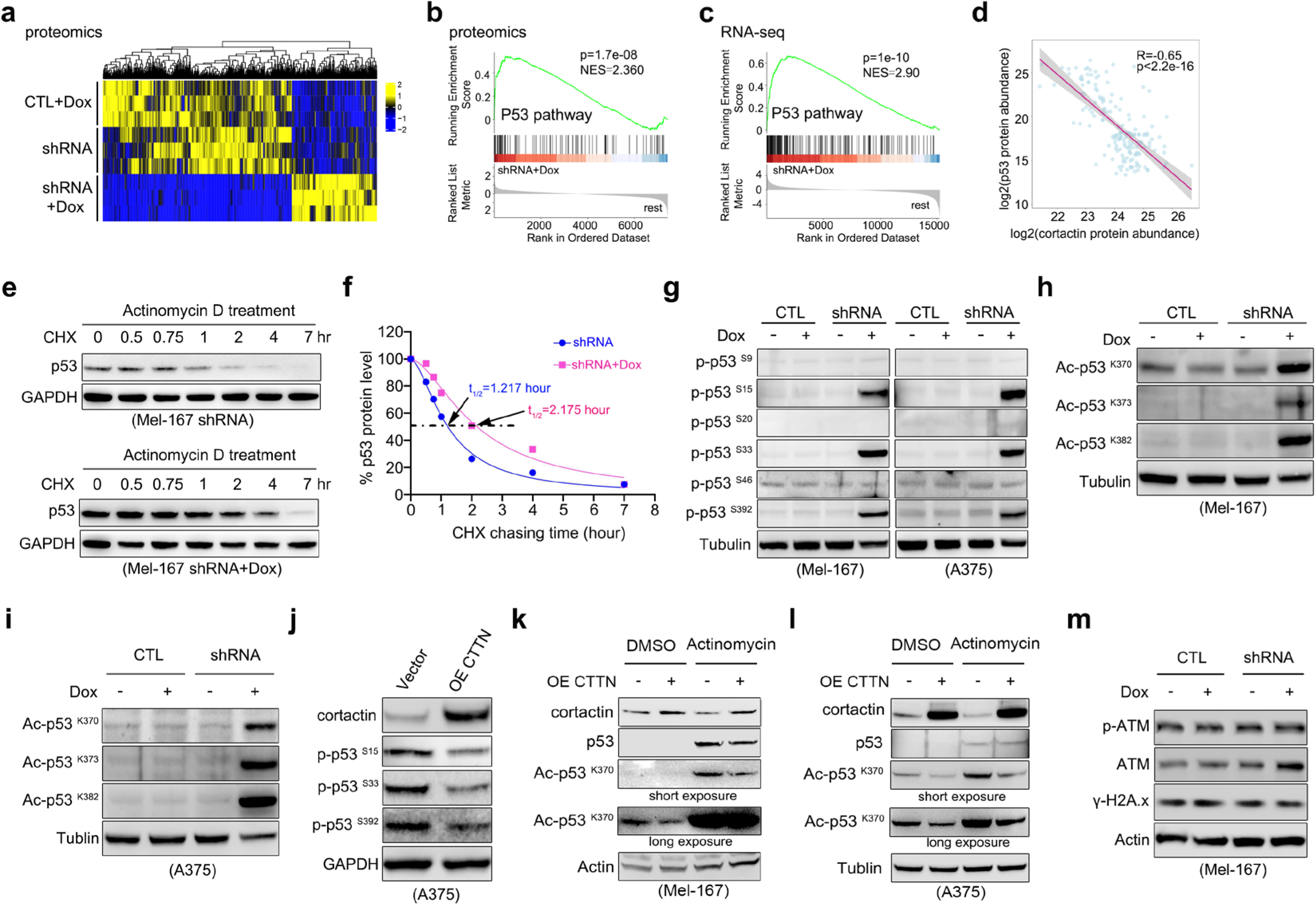
CTTN regulates p53 post-translational modifications. **a**, Heatmap depicts the proteomic changes in Mel-167 CTC with or without CTTN KD. **b,** Gene set enrichment analysis (GSEA) showed the upregulation of the p53 pathway in the CTTN KD group compared with the control group based on proteomic data. **c,** GSEA showed the upregulation of the p53 pathway in the CTTN KD group compared with the control group based on bulk RNA-seq data. **d,** Scatter plot showing the correlation between the abundance of p53 and cortactin at the protein level using public melanoma proteomic dataset from Lazaro et al cohort (PMID: 34323403). *P* value was calculated by Pearson correlation. **e**, Analysis of p53 protein stability by the cycloheximide (CHX) chase assay. p53 protein level was determined by immunoblotting. **f**, Quantification of the relative p53 protein level in the CHX chase assay of (**e**). Nonlinear regression was performed by Prism 8 software, and Does-response-inhibition (variable slope) equation was chosen to calculated the values of t1/2, half-life. **g**, Immunoblotting determining the phosphorylated p53 on multiple sites of Mel-167 and A375 with or without CTTN KD. **h-i**, Immunoblotting determining the acetylated p53 on multiple sites of Mel-167 and A375 with CTTN KD. **j**, Immunoblotting showing the changes of S15, S33, and S392 phosphorylation of p53 level in A375 with or without CTTN overexpression. **k-l**, Immunoblotting showing the decrease of acetylated p53 on K370 by overexpression of cortactin in Mel-167 and A375. **m**, Immunoblotting showing the changes of phosphorylated ATM (ser1981) and γ-H2A.x (phosphorylated-H2A.x on ser139) level in Mel-167 with or without CTTN KD.

Interestingly, a strong inverse correlation between cortactin and p53 (R=-0.65; p<2.2e-16, Fig. 3d), or cortactin and p21 (R=-0.46; p<5.8e-10, Extended Data Fig. 4h) was observed at the protein level using the human melanoma proteome atlas dataset ^46^. However, such a correlation was not observed at the mRNA level using the TCGA SKCM dataset (Extended Data Fig. 4g). Consistently, the activation of the p53 pathway in CTCs with CTTN KD was not due to changes in p53 mRNA expression (Extended Data Fig. 4f).

To understand how p53 is regulated by cortactin at the protein level, we first monitored the protein stability of p53 using a cycloheximide (CHX) chasing experiment. The results showed that the p53 was remarkably stabilized in CTTN-KD CTCs as compared with the control, with the half-life of p53 protein increasing from 1.217 hours to around 2.175 hours (Fig. 3e, f). The accumulated p53 was mainly localized in the nucleus (Extended Data Fig. 4i), where it exerts transcriptional regulatory function. Strikingly, the phosphorylation level of p53 on multiple sites, including S15, S33, and S392, was sharply elevated upon CTTN KD in both Mel-167 CTCs and A375 cells (Fig. 3g). Another important post-translation modification of p53 involved in its stabilization, the lysine acetylation, was also examined. The acetylation levels on lysine 373 (K373), lysine 382 (K382), and lysine 370 (K370) were remarkably increased in cells with CTTN KD (Fig. 3h, i). The critical modifications of p53 at these sites have collectively resulted in p53 stabilization and activation, as reported previously ^47^. Consistently, overexpression of cortactin in Mel-167 or A375 appeared to reduce the phosphorylation level of p53 at S15, S33 and S392 (Fig. 3j), and acetylation level of p53 K370, in the condition of actinomycin treatment to stabilize p53 (Fig. 3k,l). Intriguingly, the activation of p53 did not seem to be a consequence of DNA damage response as p-ATM, p-ATR and γ-H2A.x were not increased upon CTTN KD (Fig. 3m and Extended Data Fig. 7a).

These data have provided compelling evidence that cortactin regulated the stability and activity of p53 through post-translational modifications on multiple phosphorylation and acetylation sites in melanoma CTCs. To further explore if p53 is a major effector of CTTN-KD induced senescence phenotypes, we carried out a double knockdown experiment of cortactin and p53 in both Mel-167 CTCs and A375 cells. The upregulation of p21 (CDKN1A), the downregulation of p-RB, Lamin B1, and Ki-67 (MKI67) in protein and or mRNA levels with CTTN KD were at least partially rescued by p53 silencing (Extended Data Fig. 5a,b and Extended Data Fig. 6c,e). The fraction of Lamin B1 low/negative Mel-167 CTCs was significantly reduced upon p53/cortactin co-depletion (Extended Data Fig. 5c). Transcriptomic analysis showed that the repressed signature of E2F targets was rescued in double knockdown cells compared with CTTN KD alone (Extended Data Fig. 6a,d), and the rescued expression of E2Fs (E2F1, E2F2, E2F8) and its target genes (including MCM7, POLA2) were validated by RT-qPCR (Extended Data Fig. 6b,e).

Consistently, the upregulation of SASP signatures was significantly suppressed by p53/CTTN double knockdown (Extended Data Fig. 5d,f). The senescence score of p53/CTTN double knockdown cells was dramatically dropped to the level that is comparable to control knockdown cells (Extended Data Fig. 5i). Importantly, the arrest in cell proliferation was partially rescued in double knockdown cells, possibly suggesting a reduced fraction of cells entering senescence (Extended Data Fig. 5g,h; Extended Data Fig. 6f,g). All these data demonstrated that cortactin regulated CTC senescence induction in a p53-dependent manner.

### Cortactin depletion induces abnormally enlarged Rab7-positive late endosomal aggregates and results in mTOR overactivation

To identify the upstream regulators of p53 activation, we screened for candidate kinases mediating the phosphorylation of the TAD domain of p53. Phosphorylation of S15, S18, S20 sites at p53 TAD domain could impair the interaction between p53 and MDM2, resulting in decreased p53 turnover ^48^, and promote the binding of p300 to p53 leading to enhanced p53 acetylation ^48^. It has been reported that RSK2, AMPK, mTOR, ATM, ATR, DNA-PK and CDK5 are responsible for phosphorylation at S15 of p53, while CDK9, CDK5, CAK/CDK7, GSK3β, p38 MAPK could phosphorylate at S33 of p53 (Fig. 4a) ^47^. Our candidate-based biochemical screening showed that the DNA damage response markers, p-ATR S428, p-ATM and γ-H2A.x did not show upregulation upon cortactin depletion (Fig. 3m and Extended Data Fig. 7a), suggesting the canonical DNA damaging pathways was not involved in the activation of p53 in this case. The AMPK inhibitor (Dorsomorphin), RSK inhibitor (BI-D1870) and CDK5 inhibitor (PNU112455A, CDK5-IN-3) all failed to block the p-p53 S15 or p53 protein accumulation in CTTN-KD cells (Extended Data Fig. 7b, c). Furthermore, we tested the abilities of known S33-phosphorylating kinases to regulate p53 activation in CTTN-KD cells. Inhibitors of p38MAPK (Doramapimod), JNK (SP600125), CDK9 (Enitociclib), GSK3β (AR-A014418) and CDK7 (LDC4297) did not attenuate the p-p53 S33 and p53 protein expression level in cells with CTTN KD (Extended Data Fig. 7d).

**Fig. 4.**
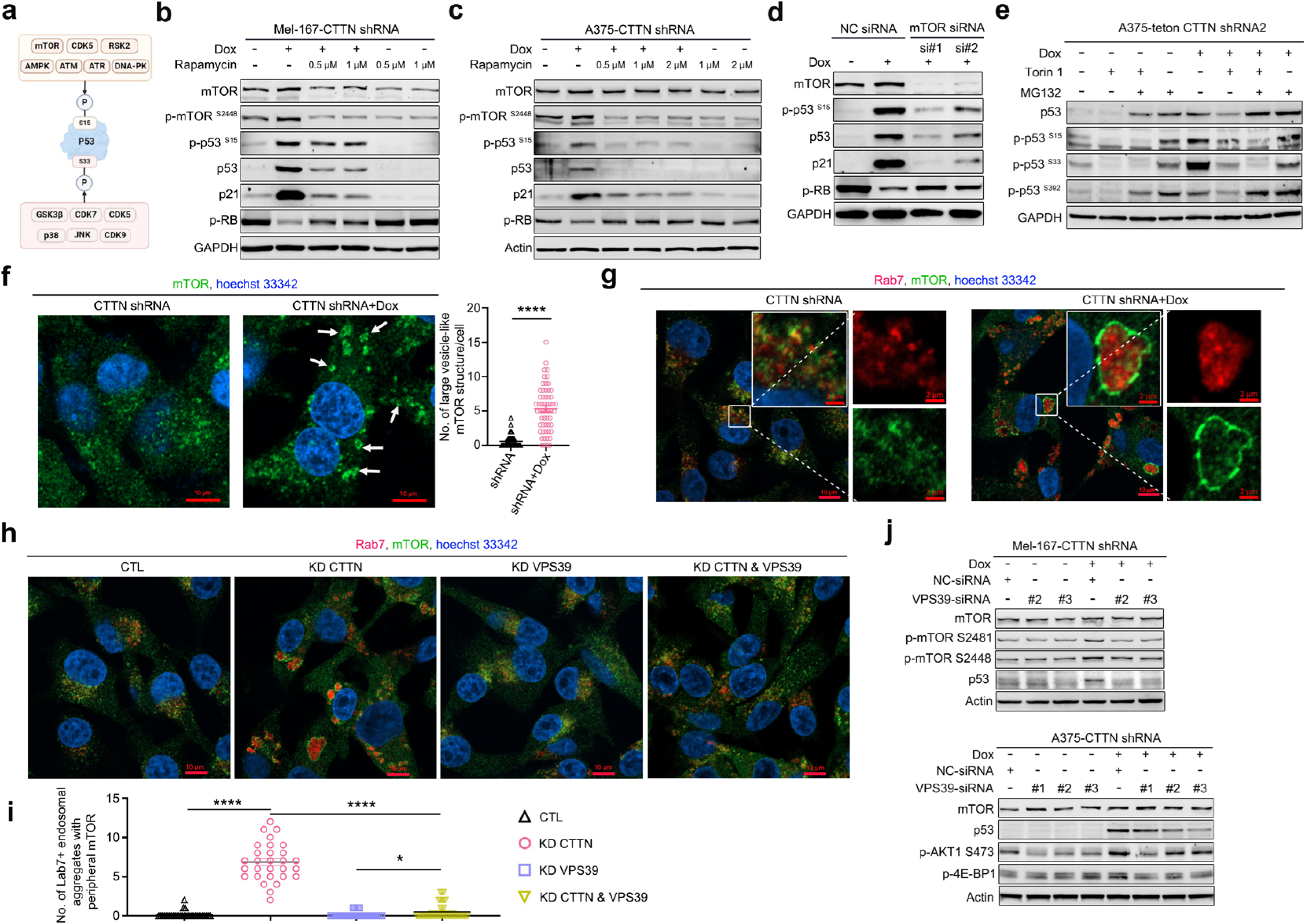
Cortactin depletion induces p53 activation through mTOR pathway activation. **a,** Picture depicted the kinases responsible for S15 and S33 phosphorylation of p53. **b,c,** Immunoblotting detection of mTOR, p-mTOR S2448, p53, p21 and phosphorylated RB S807/811 of Mel-167-teton CTTN shRNA (**b**) and A375-teton CTTN shRNA cells (**c**) with or without doxycycline and Rapamycin co-treatment for 36h. **d,** Immunoblotting detection of mTOR, p53, p-p53 S15, p21, and p-RB S807/811 upon mTOR knockdown by siRNA for 48h. **e,** Immunoblotting detection of p53, p-p53 S15, p-p53 S33, and p-p53 S392 of A375 with or without CTTN KD for 48h, followed by co-treatment of 250 nM Torin 1 and 10 μM MG132 for 4 hours. **f,** Immunofluorescence of mTOR in A375 cells with or without CTTN KD for 48h. Plot graph showing the number of large mTOR puncta (>1.5 μm) per cell. n (A375-teton CTTN shRNA)= 64 cells, n (A375-teton CTTN shRNA+Dox)= 56 cells were counted. Scale bar, 10μm. **g,** Immunofluorescence of mTOR and Rab7 in A375 cells with or without CTTN KD for 48h. Scale bar, 10μm and 2μm as indicated in images. **h,i,** Immunofluorescence of mTOR and Rab7 in A375 cells with or without CTTN and or VPS39 knockdown for 48h (**h**). Quantification of the number of enlarged vesicles (>1.5 μm) with large mTOR and Rab7 puncta (**i**). n = 32, 29, 39, 59 cell counted (from left to right). Scale bar, 10μm. **j,** Immunoblotting detection of protein levels of mTOR, p-mTOR S2448/S2481, p53, p-AKT1 S473, p-4E-BP1 in Mel-167 and A375 cells with or without CTTN and/or VPS39 knockdown for 48h. Data are represented as mean ± SEM in (**f**) and (**i**). The statistical significance was calculated by two-sided Student’s t test in (**f**) and (**i**). \**P*<0.05; **** *P*<0.0001.

The mTOR pathway was reported to be a crucial regulator of senescence, particularly in the regulation of SASP, in multiple experimental model systems ^49–51^. We further tested if treatment by the mTOR inhibitors could alleviate CTTN-KD induced p53 activation. Remarkably, Rapamycin (mTOR complex 1 inhibitor) and Torin1 (mTOR complex 1 and 2 inhibitor) treatment strongly blocked the accumulation of p53 and the phosphorylation at S15 in melanoma cells with cortactin depletion, which also led to the decreased p21 protein level and increased phosphorylated RB protein level (Fig. 4b,c and Extended Data Fig. 7e). CTTN KD resulted in increased phosphorylation of mTOR on S2448 and S2481, as well as its substrates 4E-BP1 on T37/46, AKT1 on S473 (Fig. 4b,c and Extended Data Fig. 7e). As PI3K/AKT signaling was reported to activate mTOR, we thus test if pan PI3K inhibitor could block mTOR and p53 activation. Wortmannin (pan-PI3K inhibitor) treatment could efficiently reduce the level p-mTOR S2448/S2481, p-AKT1 S473, p-4E-BP1 T37/46, p53, p-p53 S15 and p21 (Extended Data Fig. 7e). The ability of mTOR complex to activate p53 pathway is further validated using genetic depletion of mTOR, Raptor and Rictor by siRNA treatment (Fig. 4d and Extended Data Fig. 7f). Therefore, mTOR signaling was the key upstream regulator of CTTN KD-induced p53 activation and senescence induction.

Since mTOR signaling is known to promote mRNA translation efficiency ^52,53^, we quantified polysome-bound p53 mRNA level in CTTN-KD cells and found that p53 mRNA translation was not affected (Extended Data Fig. 7g). Further, we used MG132 to block p53 degradation in the present of Torin1 treatment and found that the decreased p-p53 S15 caused by mTOR inhibition was not a sequential consequence of the decrease of p53 (Fig. 4e), suggesting mTOR complex can phosphorylate p53 at S15. Interestingly, the p-p53 S33, but not S392, also kept in a low level in cells with both Torin 1 and MG132 treatment regardless the high level of p53 it contained (Fig. 4e), indicating that S33 of p53 may be a novel phosphorylation site of mTOR complex in our experimental system. Functionally, blocking mTOR signaling using mTOR inhibitors consistently reduced the expression level of SASP signature genes (Extended Data Fig. 7h) and the β-gal positivity in CTTN-KD cells (Extended Data Fig. 7i).

mTOR is reported to be active at the plasma membrane, outer mitochondrial membrane, endosomal vesicles, lysosome, cytosol, nucleus and ribosome ^54–57^. Strikingly, immunocytochemistry showed that CTTN KD induced the formation of large mTOR ring-like structures that were larger than 1.5 μm in diameter (Fig.4f). To investigate which types of organelles could co-localize with these mTOR-enriched structures, cells were co-stained with mTOR and late endosome marker (Rab7) or early endosome marker (Rab5) or lysosome marker (LAMP2). CTTN KD did not change the expression level of Rab5 and LAMP2 (Extended Data Fig. 8a) and the mTOR-enriched structures neither co-localize with early endosome nor lysosome (Extended Data Fig. 8b, d), although large Rab5 positive puncta (Extended Data Fig. 8b, c) and a dispersive distribution pattern of lysosome was observed in CTTN KD cells (Extended Data Fig. 8d, e). CTTN KD induced dramatic aggregation of Rab7 positive late endosomes (Extended Data Fig. 8f,g). Intriguingly, we found that those aberrant ring-like mTOR structures were co-stained with late endosomal structures (Fig.4g), with the strongest mTOR signal detected along the peripheral edges (Extended Data Fig. 8h). These appeared to be mTOR-enriched Rab7^+^ late endosomal aggregates. Consistently, cortactin was also found to be co-localized with late endosome in A375 cells overexpressing HA-cortactin (Extended Data Fig. 8i,j), suggesting its potential involvement in the regulation of late endosome tethering and homeostasis. Importantly, silencing the expression of VPS39, a component of HOPS complex, which is required for homotypic tethering, fusion and maturation of late endosomes ^58^, significantly impaired the formation of large mTOR^+^ Rab7^+^ late endosomal aggregates (Fig.4h,i), and inhibited mTOR overactivation and p53 activation (Fig. 4j). We speculated that blocking the maturation from early endosome to late endosome may reduce the formation of late endosomal aggregations and downregulate mTOR/p53 signaling. Indeed, depletion of RMC1 (encoding C18orf8), a component of Mon1-Ccz1 complex mediating Rab5-to-Rab7 conversion ^59,60^, resulted in decreased mTOR/p53/p21 signaling (Extended Data Fig. 8k). Interestingly, the knockdown of VPS8, a component of CORVET complex mediating the tethering of early endosomes ^60^, and the depletion of ARL8A/B, which mediate the tethering between late endosome and lysosome ^61^, both of which did not affect the activation of p53 (Extended Data Fig. 8l,m), suggesting that likely CTTN KD induced the defects in the tethering and homeostasis within late endosomes. These data suggested that the large Rab7-positive late endosomal aggregates in CTTN-KD cells likely served as a “molecular trap” for mTOR accumulation and overactivation.

Taken together, by localizing to Rab7 positive endosomes, cortactin was required for controlling the late endosomal tethering and homeostasis that was critical for mTOR activation. CTTN KD resulted in aberrant aggregations of Rab7-postive late endosomes, where the mTOR protein complex accumulated and became activated, subsequently leading to S15 and S33 phosphorylation and p53 activation.

### Cortactin regulates mitochondrial functionality and ROS homeostasis

Mitochondrial dysfunction is considered a hallmark of cellular senescence and aging^40,62,63^. Our GSEA and signature heatmap analyses showed that the expression of ROS pathway gene set and oxidative phosphorylation (OXPHOS) gene set were positively enriched in CTTN-KD group as compared to the control (Fig. 5a, b and Extended Data Fig. 9a-c). Given that the ROS signature was one of the top enriched pathways in CTTN-KD CTCs (Extended Data Fig. 9d), we hypothesized that the senescence induction in our experimental settings may be associated with mitochondrial dysfunction. We first used a transmission electron microscope (TEM) to capture the ultrastructural changes of mitochondria in Mel-167 and A375 cells. CTTN-KD CTCs and A375 demonstrated significantly enlarged and swollen mitochondrial morphology (Fig. 5c,d), reduced cristae length, cristae number per mitochondria and the ratio of cristae length to outer mitochondria membrane, as compared to control cells (Fig. 5e-f,h-j). Furthermore, the mitochondrial DNA (Fig. 5k and Extended Data Fig. 9e,f) and mRNA expression (Fig. 5l) was elevated upon CTTN KD. Using the mitoSOX Red dye, we found the mitochondrial superoxide level was dramatically increased (Fig. 5m,n and Extended Data Fig. 9g,h), and mitochondria membrane potential was also remarkably decreased in CTTN-KD CTCs (Fig. 5o). These data collectively suggested cortactin is a critical regulator of mitochondrial structure, morphology, oxidative function and ROS generation in melanoma CTCs. The dysfunctional mitochondria and ROS accumulation in CTCs could contribute significantly to senescence induction.

**Fig. 5.**
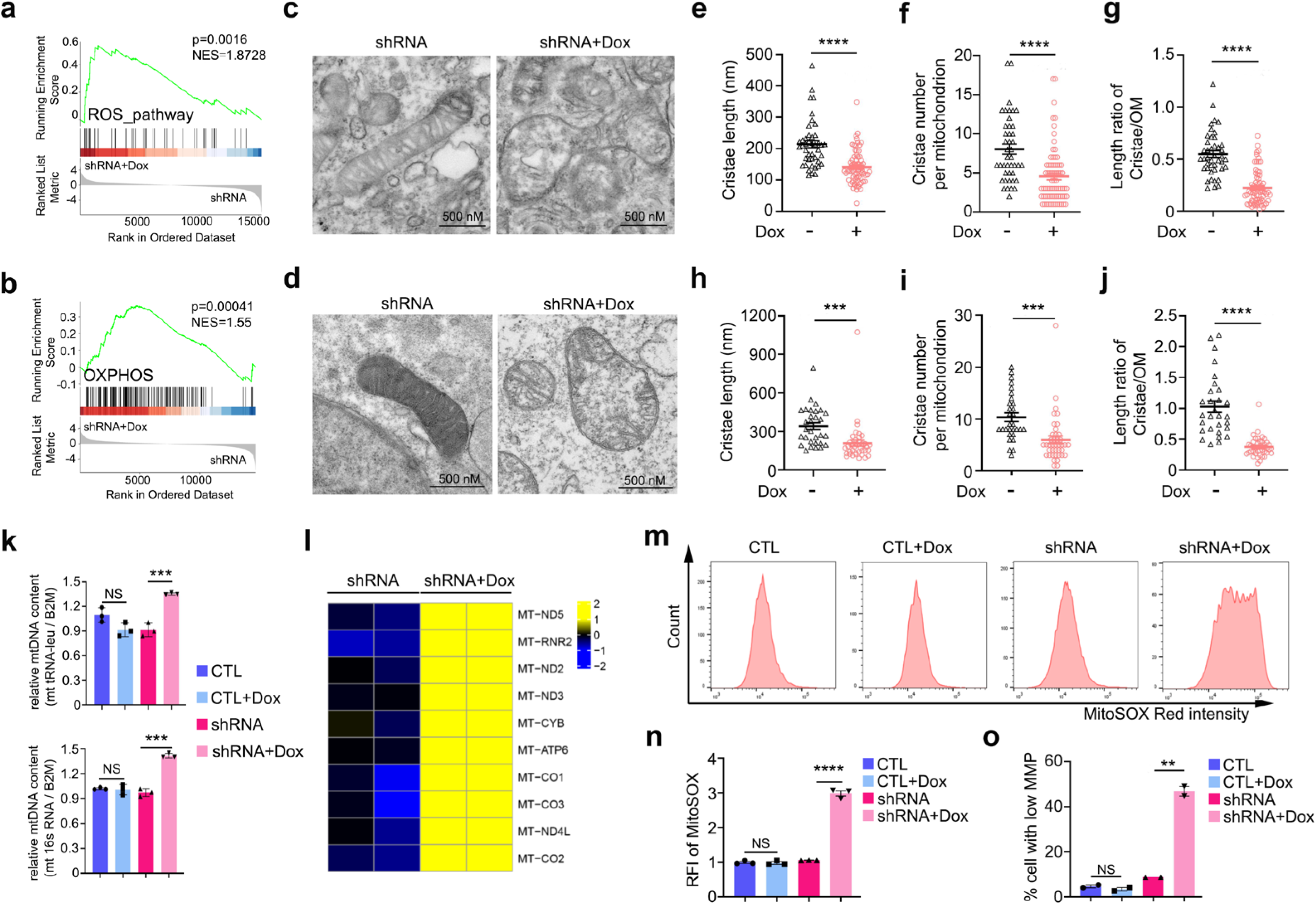
Cortactin depletion leads to mitochondrial dysfunction and ROS generation. **a,b**, GSEA of ROS pathway (**a**) and OXPHOS gene set (**b**) comparing Mel-167 with or without CTTN KD by 4 days’ doxycycline induction. **c,d**, Mitochondrial ultrastructure of Mel-167 (**c**) and A375 (**d**) with or without CTTN KD by 3 days’ doxycycline induction, images were captured by TEM. **e-g**, Scatter plot showed the quantification of cristae length (**e**), cristae number per mitochondria (**f**) and length ratio of cristae/ outer mitochondrial membrane (**g**) of Mel-167 cells (**c**). n of (**e-g**)= 43 and 72 mitochondria counted for cells with or without CTTN KD, respectively. **h-j,** Scatter plot showed the quantification of cristae length (**h**), cristae number per mitochondria (**i**) and length ratio of cristae/ outer mitochondria membrane (**j**) of A375 cells (**d**). n of (**h-j**)= 30 and 40 mitochondria counted for cells with or without CTTN KD, respectively. **k**, Relative mitochondrial DNA content determined by qPCR, by normalizing mt tRNA-leu or mt 16s rRNA DNA content to nuclear B2M DNA content of Mel-167 cell line. n=3 independent experiments. **l**, Heatmap showed the upregulated mRNA encoded by mitochondria of Mel-167 cells with or without CTTN KD for 3 days of doxycycline treatment. **m, n**, Mitochondrial superoxide of Mel-167 cells determined by MitoSOX red staining followed by flow cytometry analysis (**m**). Bar graph (**n**) shows the quantification of mitochondrial superoxide. Doxycycline treatment for 3 days. n=3 independent experiments. **o**, Mitochondrial membrane potential analyzed by JC-1 staining followed by flow cytometry. n=2 independent experiments. Data are represented as mean ± SEM in (**e-g**), (**h-k**), (**n**) and (**o**). The statistical significance was calculated by two-sided Student’s t test in (**e-g**), (**h-k**), (**n**) and (**o**). \*\**P*<0.01; \*\*\**P*<0.001; **** *P*<0.0001.

### p53 activation and mitochondrial ROS form a signal amplification loop driving stable senescence

ROS has been widely reported to activate p53, and p53 in turn regulates cellular redox status through various distinct mechanisms ^64,65^. To dissect the functional relationship between p53 activity and ROS regulations, we performed a time-course analysis in CTCs with inducible CTTN KD and monitored p53 protein level, cell cycle, and mtROS levels over time. We found that p53 began to accumulate at 24h post doxycycline treatment, then dramatically increased at 48 and 72h post doxycycline treatment (Fig. 6a, Extended Data Fig. 10a), which coincided with G0/G1 arrest (Extended Data Fig. 10b,c). Interestingly, mtROS did not increase 24h post doxycycline treatment (Extended Data Fig. 10d,f), but surged at 48h (Extended Data Fig. 10e,f). Time-course quantification of p53 protein, cell cycle status, and mtROS level in both Mel-167 CTC and A375 cells with doxycycline treatment for 0h, 24h, 48h and 72h revealed that mtROS accumulation occurred lagging behind p53 activation and G0/G1 arrest (Fig. 6b and Extended Data Fig. 10g), implicating mitochondrial dysfunction may be a consequence of p53 activation. The striking elevation of p53 abundance from 48-72 hours post doxycycline treatment, correlated with a sharp rise in mtROS accumulation at the same time scale, suggesting a possible positive crosstalk between these two biological processes (Fig. 6b and Extended Data Fig. 10g).

**Fig. 6.**
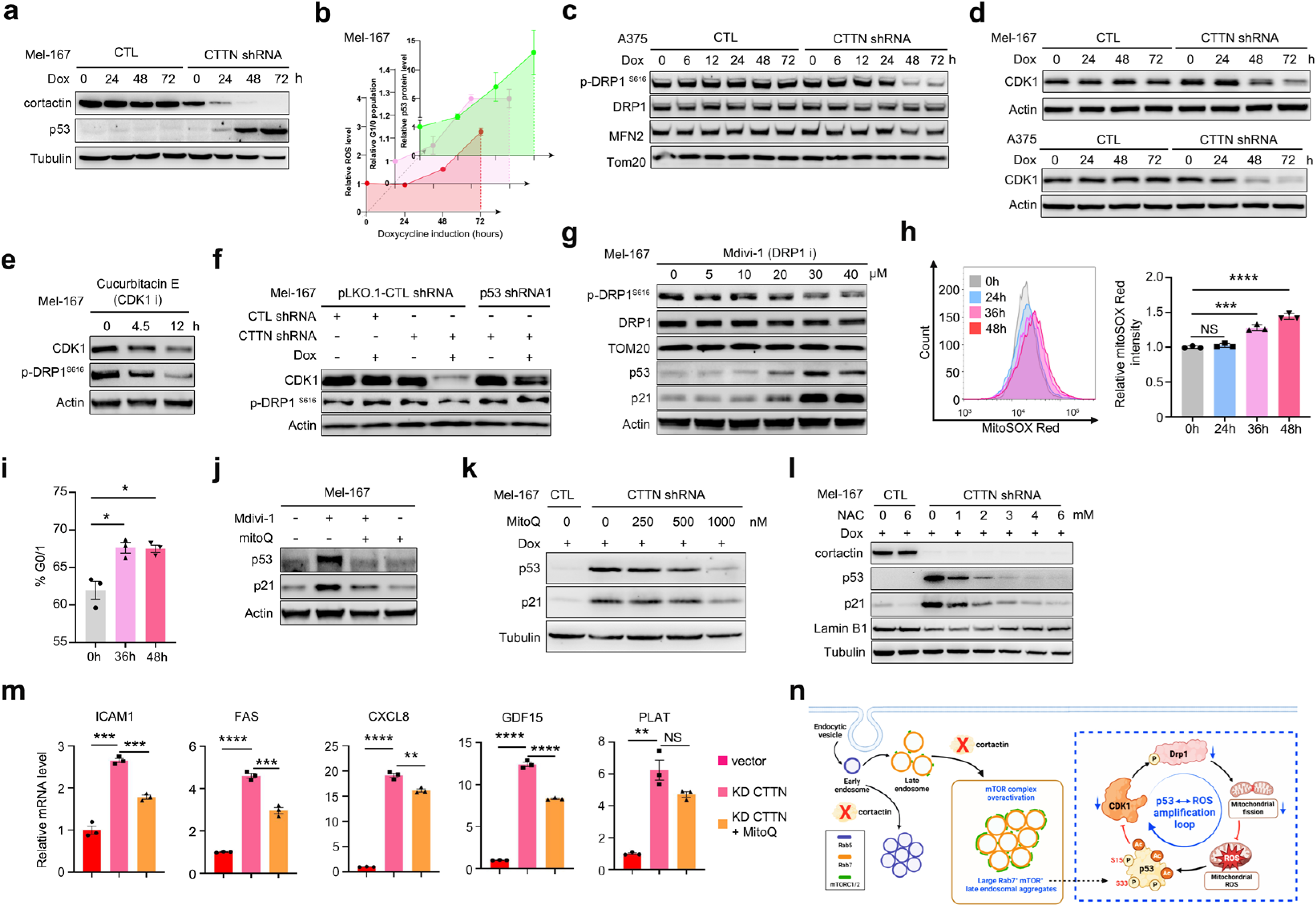
p53-mtROS positive feedback regulation forms a signal amplification loop driving senescence induction and maintenance. **a,** Immunoblotting of cortactin and p53 protein level of Mel-167 with or without CTTN KD for the indicated time by doxycycline treatment. **b**, Line chart depicted the relative mtROS level, relative G0/G1 population, and p53 protein level of Mel-167 cells with or without CTTN KD by doxycycline induction for 0h, 24h, 48h, and 72h. n=3 independent experiments. **c**, Immunoblotting showed p-DRP1 S616, DRP1, MFN2 of A375 with or without doxycycline induction for the indicated time. **d**, Protein level of CDK1 of Mel-167 and A375 with or without doxycycline induction for indicated time to knockdown cortactin. **e,** CDK1 and p-DRP1 S616 protein level of Mel-167 cells in the presence of 12h treatment of 10 uM CDK1 inhibitor, Cucurbitacin E. **f**, Immunoblotting showed p53, CDK1, p-DRP1 S616 protein level of Mel-167 with or without cortactin and or p53 knockdown. Cortactin shRNA was cloned into the teton-pLKO.1-Puro vector while p53 shRNA was cloned into pLKO.1-Blast vector. **g**, Protein level of p-DRP1 ser616, p53, and p21 of Mel-167 in the presence of Mdivi-1 treatment for 36h. **h**, Relative mtROS level of Mel-167 with or without 10 uM Mdivi-1 treatment determined by mitoSOX Red staining. n=3 independent experiments. **i**, Cell cycle analysis of G0/1 phase of Mel-167 cells in the presence of 10 μM Mdivi-1 treatment for the indicated time. n=3 independent experiments. **j**, Protein level of p53 and p21 of Mel-167 in the presence of 10 μM Mdivi-1 and/or 1000 nM MitoQ treatment. **k,l**, p53, p21, and or Lamin B1 protein level of Mel-167 pretreated with or without doxycycline to knockdown cortactin for 24h then treated with MitoQ (**k**) or NAC (**l**) treatment for 48h. **m,** RT-qPCR determination of ICAM1, FAS, CXCL8, GDF15, and PLAT expression of Mel-167 with or without Cortactin knockdown and 1000 nM MitoQ treatment. n=3 independent experiments. **n**, Picture illustrated the amplification loop of p53 accumulation and mtROS elevation. Data are represented as mean ± SEM in (**b**), (**h**), (**i**) and (**m**). The statistical significance was calculated by two-sided Student’s t test in (**h**), (**i**) and (**m**). \**P*<0.05; \*\**P*<0.01; \*\*\**P*<0.001; **** *P*<0.0001; NS, not significant.

The dynamic processes of mitochondrial fission and fusion are tightly regulated, determining mitochondrial shape, and influence mitochondrial function and turnover ^66^. For example, mitochondrial fission and fusion mediate energy output, ROS production, and mitochondrial quality control ^67^. The fission of mitochondria is positively regulated by FIS1, MFF, and DRP1, while the fusion of mitochondria is positively regulated by Opa1, MFN2, MFN1, and OPA1 ^68^. To understand the detailed mechanisms of mitochondrial dysfunction induced by p53 activation, the regulators of mitochondrial fission and fusion dynamics, including DRP1 S616 phosphorylation and MFN2, were assessed by western blot analysis. DRP1 S616 phosphorylation was significantly decreased in CTTN-KD cells 48 hours after Dox treatment (Fig. 6c, and Extended data fig. 11a), indicating the mitochondrial fission was largely impaired. CDK1, PKCd, ERK1/2, and CaMKII have been shown to phosphorylate DRP1 at S166 directly ^68^. Among these kinases, CDK1 was found to be strongly suppressed in CTTN-KD cells (Fig. 6d), presumably due to p53 activation ^45,69^. We thus hypothesized that the reduced DRP1 phosphorylation might be due to the suppression of CDK1 expression in CTTN-KD cells. Interestingly, inhibition of CDK1 by Cucurbitacin E could markedly block the phosphorylation of DRP1 at S616 (Fig. 6e and Extended Data Fig. 11b). These data suggested that p53 activation in CTTN-KD cells could significantly impair the phosphorylation of DRP1 by inhibiting CDK1 expression. Indeed, CTTN KD markedly inhibited CDK1 mRNA expression in a p53-dependent manner (Extended Data Fig. 11c, d). p53 silencing also partially rescued the p-DRP1 S616 and CDK1 protein levels (Fig. 6f and Extended Data Fig. 11e). Furthermore, treatment by DRP1 specific inhibitor Mdivi-1 was sufficient to activate p51/p21 (Fig. 6g and Extended data fig. 11h,i), accompanied by increased mtROS (Fig. 6h and Extended data fig. 11f,g) and G0/G1 arrest (Fig. 6i and Extended Data Fig. 11j) in both Mel-167 CTCs and A375 cells. Consistently, DRP1 KD by shRNA also led to G0/G1 arrest in A375 cells (Extended Data Fig. 11k). Therefore, the activation of p53 by mTOR resulted in reduced CDK1 expression and impaired DRP1 phosphorylation at S616, inducing mitochondrial fission defects and mtROS accumulation.

To test if elevated mtROS production may feedback to enhance p53 activation, mtROS scavenger MitoQ was applied to Mdivi-1 treated cells. Interestingly, p53/p21 activation was partially inhibited (Fig. 6j and Extended data fig. 11l). The ability of MitoQ to rescue p53/p21 elevation was also evident in CTTN-KD cells (Fig. 6k, Extended data fig. 11n). Similarly, a partial rescue was also observed using other anti-oxidants, such as NAC (N-Acetyl-D-cysteine) treatment in Mel-167 CTCs (Fig. 6l, Extended data fig. 11m), as well as Tempol (a superoxide dismutase mimic) treatment in A375 cells (Extended data fig. 11o). ROS clearance by MitoQ treatment in CTTN-KD cells also significantly suppressed SASP upregulation including ICAM1, FAS, CXCL8, GDF15, and PLAT (Fig. 6m), and substantially impaired β-gal formation (Extended data fig. 11p).

Taken together, the initial p53 activation by the aberrant mTOR overactivation was subsequently enhanced and amplified by the “p53 ---| CDK1→p-DRP1 ---| mtROS→ p53” loop, leading to prolonged p53 activities and stable senescence maintenance in CTTN-KD CTCs (model outlined in Fig. 6n).

### A “One-two punch” sequential therapy involving CTTN KD followed by senolytic drug treatment enables effective CTC clearance

Senescent cells often exhibit increased levels of anti-apoptotic BCL2 family proteins to protect cells from apoptosis ^70^. The senolytic drugs, Navitoclax (also known as ABT-263) and ABT-737, selective inhibitors of Bcl-2, Bcl-xL, and Bcl-W, were reported to effectively eliminate senescent tumor cells by triggering apoptosis ^71^. These promising observations motivated us to test a novel “One-two punch” therapeutic strategy, involving sequential treatment of CTCs with CTTN KD followed by senolytic drug treatment to completely eliminate CTCs and metastatic dormant clones.

Remarkably, BCL2L1 (Bcl-xL) mRNA was significantly increased in Mel-167 CTCs and xenograft tumors with CTTN KD (Fig. 7a,b). Mel-167 CTCs with CTTN KD was much more sensitive to ABT-263 and ABT-737 treatment than the control group (Fig. 7c,d). ABT-263 and ABT-737 treatment induced strong apoptosis in CTTN-KD CTCs, as indicated by decreased Bcl-xL, pro-caspase 9, pro-caspase 8, pro-caspase 3, full length PARP and the increased cleaved-caspase 3, cleaved-PARP with a drug concentration down to 0.195 μM. In contrast, CTTN KD alone conferred minimal activation of caspase 3 (Fig. 7e). Sequential therapy using CTTN KD followed by low dosage of Bcl2 inhibitor treatment (30 mg/Kg/day) led to the remarkable regression of Mel-167 tumorigenic growth, with the tumor size and weight significantly shrinked as compared to the control group, CTTN-KD group, or ABT-263 treatment alone (Fig. 7f-h). Consistently, tumor tissues with the sequential therapy showed much more apoptotic cell death than the other group (Fig. 7i, j).

**Fig. 7.**
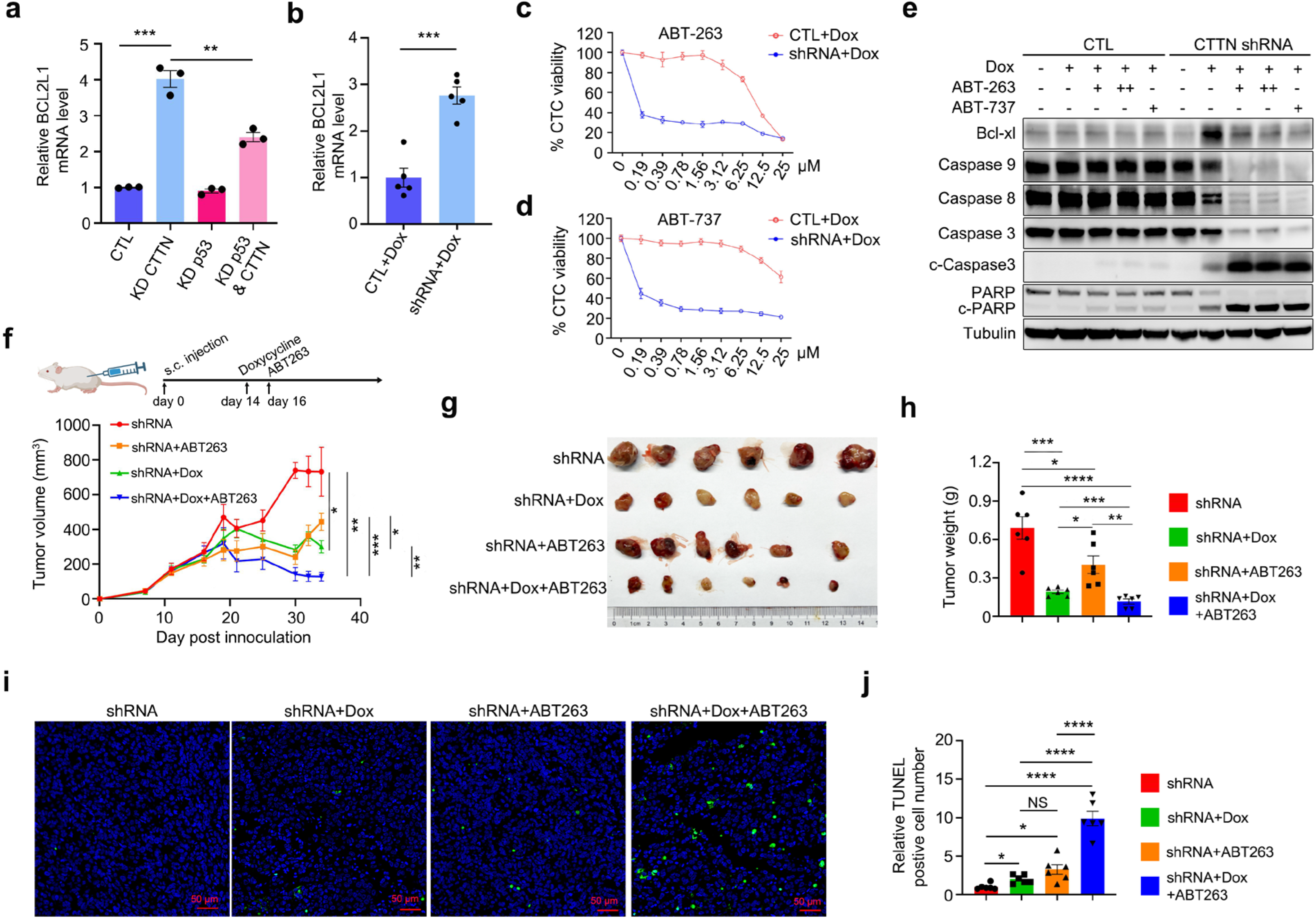
Cortactin depletion-induced senescent CTCs are susceptible to apoptotic cell death upon anti-BCL2 senolytic treatment. **a**, mRNA level of BCL2L1 (Bcl-xl) of Mel-167 cell with or without cortactin and or p53 knockdown, determined by RT-qPCR. n=3 independent experiments. **b**, mRNA level of BCL2L1 of xenografts established by s.c injection of Mel-167 to NCG mice. n=5 tumors. **c,d**, Cell viability of Mel-167 cells with or without cortactin knockdown treated by ABT-263 (**c**) and ABT-737 (**d**). Cells were pretreated with 300 ng/ml doxycycline to induce cortactin knockdown for 2 days, then cells were treated with ABT263 or ABT737 for 1 day. n=3 independent experiments. **e**, Immunoblotting of Bcl-xl and indicated apoptotic markers of Mel-167 with or without CTTN KD in the presence of ABT-263 (0.195 μM or 1.56 μM) or ABT-737 (1.56 μM). **f**, Tumor growth curve of Mel-167 xenografts established through s.c. injection. Doxycycline, 1mg/ml in drinking water. ABT263, 30 mg/kg/day. n=6 mice. **g**, Picture showed the tumors collected on the experiment endpoint of (**f**). **h**, Bar graph showed the tumor weight of (**g**). n=6 tumors. **i**, Representative images of TUNEL staining of the apoptotic cells of xenografts from (**g**). **j**, Quantification of TUNEL positive cell of (**i**). n=6 tumors. Data are represented as mean ± SEM in (**a**-**d**), (**f**), (**h**), (**j**). The statistical significance was calculated by two-sided Student’s t test in (**a**), (**b**), (**f**), (**h**), (**j**). \**P*<0.05; \*\**P*<0.01; \*\*\**P*<0.001; **** *P*<0.0001.

These preclinical data demonstrated the potential of using CTTN KD followed by senolytic drugs to effectively induce CTC apoptosis from senescent state during cancer metastatic transition, which may open a new avenue for anti-metastasis therapies.

## Discussion

Both quiescence and senescence have been suggested to play a key role in the formation of metastatic dormancy, which typically occurs in disseminated tumor cells (DTCs) that are proliferation-arrested in distant organs for a period of time before resuming their growth upon appropriate stimulus within the microenvironment ^1,3,4^. This state is characterized by the arrest of cell division, balancing proliferation with cell death, and maintaining survival without expansion ^72^. The biological complexity of metastatic dormancy lies in its regulation by intrinsic factors within the tumor cells and extrinsic signals from the surrounding microenvironment. Intrinsically, dormancy is influenced by genetic and epigenetic changes that affect cell cycle regulators, such as p21, p27, and signaling pathways like ERK/p38 MAPK ^73^. Extrinsic factors include immune surveillance, extracellular matrix (ECM) interactions, angiogenic suppression, and signals from stromal cells that collectively create a non-permissive environment for dormant niche formation in metastatic sites ^74^. Thus, a comprehensive understanding of the mechanisms mediating metastatic dormancy and reactivation is crucial for devising effective strategies to eliminate these life-threatening metastatic seeds.

Our study revealed for the first time that metastatic dormancy may actively take place in CTCs transiting in blood, before they further disseminate into distant organs to become DTCs. While the conventional thought is that most CTCs will undergo anoikis and become apoptotic in the bloodstream due to shear flow forces and various environmental stressors ^75–77^, our work provided substantial evidence that a delicate genetic circuit mediated by the “Cortactin-mTOR-p53-mitochondria”axis regulates metastatic senescence and dormancy of CTCs during blood-borne cancer dissemination.

The actin cytoskeleton regulator, cortactin, was found to play a critical role in the senescence induction of CTCs. Cortactin was originally discovered as a protein substrate of the Src kinase that regulates actin polymerization and cytoskeleton dynamics ^78,79^. It is reported to be critically involved in the formation of lamellipodia and membrane ruffles that are key subcellular structures supporting tumor cell migration and invasion ^80,81^. Using patient-derived CTC model systems, we reported an unexpected role of cortactin in regulating metastatic senescence and dormancy of CTCs. Upon cortactin depletion, these CTCs can enter a unique senescence state induced by mTOR pathway overactivation (Fig. 4b,c and Extended Data Fig. 7e). Mechanistically, CTTN-KD cells exhibited abnormal late endosomal tethering, forming unusually enlarged Rab7-postive late endosomal aggregates, on the periphery of which mTOR accumulated and activated (Fig. 4g, Extended Data Fig. 8h). The overactivation of mTOR complex on these aberrant vesicular structures resulted in p53 phosphorylation and stabilization, leading to robust senescence induction. The initial p53 activation significantly suppressed CDK1 expression, leading to the inhibition of DRP1 phosphorylation and mitochondrial fission dynamics in CTTN-KD CTCs (Fig. 6c-f). Compromised mitochondrial fission and the subsequent dysfunction, characterized by the abnormal mitochondrial morphology and structure (Fig. 5c,d), increased mitochondrial DNA (Fig. 5k), declined mitochondrial membrane potential (Fig. 5o), and ROS accumulation (Fig. 5m,n), further amplified p53 activation and thus maintained CTCs in a stable senescence state, that were cell-cycle arrested, apoptosis-resistant, and metabolically active by secreting abundant SASP factors (Fig. 6n). Hence, our work discovered a crucial signal amplification mechanism driven by the“p53 ---| CDK1→ p-DRP1 ---| mtROS→p53”positive feedback loop, leading to prolonged p53 activation and stable senescence maintenance in CTTN-KD CTCs.

The remarkable heterogeneity and plasticity of CTCs exhibiting either proliferative or pre-senescent states may maximize their survival and endurance in circulation. Indeed, single-cell RNA seq of early cultures of melanoma CTCs demonstrated heterogenous expression of CTTN within individual CTCs (Extended Data Fig. 2a,b). The expression level of CTTN exhibited an inverse correlation with various senescence hallmarks (Extended Data Fig. 2c,d). It can be speculated that CTTN ^high^ CTCs may possess a high proliferative potential and can extravasate and establish distant metastasis more efficiently than CTTN ^low^ CTCs, and they might be actively targeted by immune cells and sensitive to drugs acting on proliferating tumors; In contrast, CTTN ^low^ CTCs may be rapidly primed into a state of dormancy upon dissemination, potentially shielding them from immunological surveillance and other insults. Given the ability of senescent CTCs to secrete environment-modifying SASPs, they may slowly promote vascular remodeling until they are ready for extravasation. Therefore, it is foreseeable that similar to DTCs, senescent CTCs may serve as critical reservoirs for disease recurrence and therapeutic resistance, emphasizing the importance of detecting, characterizing, and targeting this unique population to prevent metastatic spread ^7,75^.

Based on these mechanistic insights, we proposed a sequential “One-two punch” therapeutic regimen using CTTN depletion followed by the senolytic drugs to eliminate viable CTCs. Disseminating CTCs with high CTTN expression may possess higher proliferative capacity and metastatic potential. Therefore, they can be firstly targeted for senescence induction (e.g. by blocking CTTN expression), leading to proliferation arrest and possibly a transient slowdown of the metastatic progression. Once CTCs were induced to enter a relatively stable senescence state, they had to maintain cell viability by upregulating anti-apoptotic signatures, thus creating a molecular dependency on these genes (Fig. 7a,b). Using senolytic drugs targeting these anti-apoptotic genes such as BCL2L1, massive apoptotic cell death was induced (Fig. 7e), leading to robust inhibition of CTC growth *in vitro* and metastatic tumor growth *in vivo* (Fig. 7c-j). Such proof-of-concept preclinical study may provide a framework for future clinical development of sequential therapies targeting blood-borne metastasis in melanoma.

This work suggested mTOR/p53 pathways are key effectors downstream of cortactin, molecularly reminiscent of oncogene-induced senescence (OIS) ^16,18,82^. Different oncogenic stimuli can induce senescence with distinct mechanisms, including Ras^V12^, BRAF^V600E^, or PI3K/mTOR in various cancer types. Nevertheless, the activation of the downstream p53/p21 axis or p16 appears essential ^83^ . It has been reported that PTEN loss or PI3K/mTOR activation-driven senescence occurs without apparent DNA damage response, which activates p53 through phosphorylation of S15 ^84^. This is consistent with our observation that the canonical DNA damage pathway activation was not evident in our experimental CTC models (Fig. 3m and Extended Data Fig. 7a), despite a remarkable accumulation of mtROS in CTTN-KD CTCs (Fig. 6b and Extended Data Fig. 10g). Intriguingly, some studies have suggested a role of PI3K/mTOR signaling in overcoming OIS. For example, Vredeveld et al reported that the PI3K activation can overcome BRAF^V600E^-induced senescence and contribute to melanoma progression from nevi ^85^. Another report has shown that simultaneous Cdkn2a and Lkb1 inactivation results in mTORC1/2 activation and overcomes BRAF^V600E^-induced growth arrest in a transgenic mouse model of melanoma development ^86^. On the contrary, our study demonstrated that the endosomal activation of mTOR represented a key mechanism promoting senescence induction in melanoma CTCs. Genetic depletion using siRNAs against the mTOR complex (mTOR, Raptor, Rictor), have consistently shown to alleviate p53/p21 activation in CTTN-KD cells (Fig. 4d and Extended Data Fig. 7f). Similarly, pharmacological inhibition by mTOR inhibitors significantly inhibited p53/p21 activation (Fig. 4b,c and Extended Data Fig. 7e), SASP expression (Extended Data Fig. 7h), and beta-gal formation (Extended Data Fig. 7i). Therefore, in contrast to the previous reports about the oncogenic role of PI3K/mTOR in overcoming OIS during melanocytic nevi transformation, our work suggested an essential role of mTOR activation in CTC senescence induction during metastatic progression. Thus, there may be context-dependent roles of PI3K/mTOR signaling in OIS, which could either promote or suppress senescence depending on cancer types or stages of cancer progression, providing a broad clinical implication when therapeutic strategies are being developed to treat metastasis.

In summary, our study has provided substantial evidence that metastatic dormancy could occur in transiting CTCs within the blood circulation before their arrival in distant organs as DTCs. The “Cortactin-mTOR-p53-mitochondria” regulatory network was critically involved in senescence induction of CTCs. Further mechanistic investigation is undoubtedly needed to elucidate how cortactin regulates Rab7-positive late endosome complex and function. Nonetheless, our study highlights a potential survival strategy employed by a subset of CTCs with low CTTN expression. These cells may be primed to enter a senescent and dormant state within the hostile blood microenvironment, enabling them to persist prior to extravasation and metastatic outgrowth. A “One-two punch” treatment strategy using CTTN KD followed by senolytic therapies may offer a powerful approach to suppress CTC-mediated metastatic progression (Figure 8, Working Model).

**Fig. 8.**
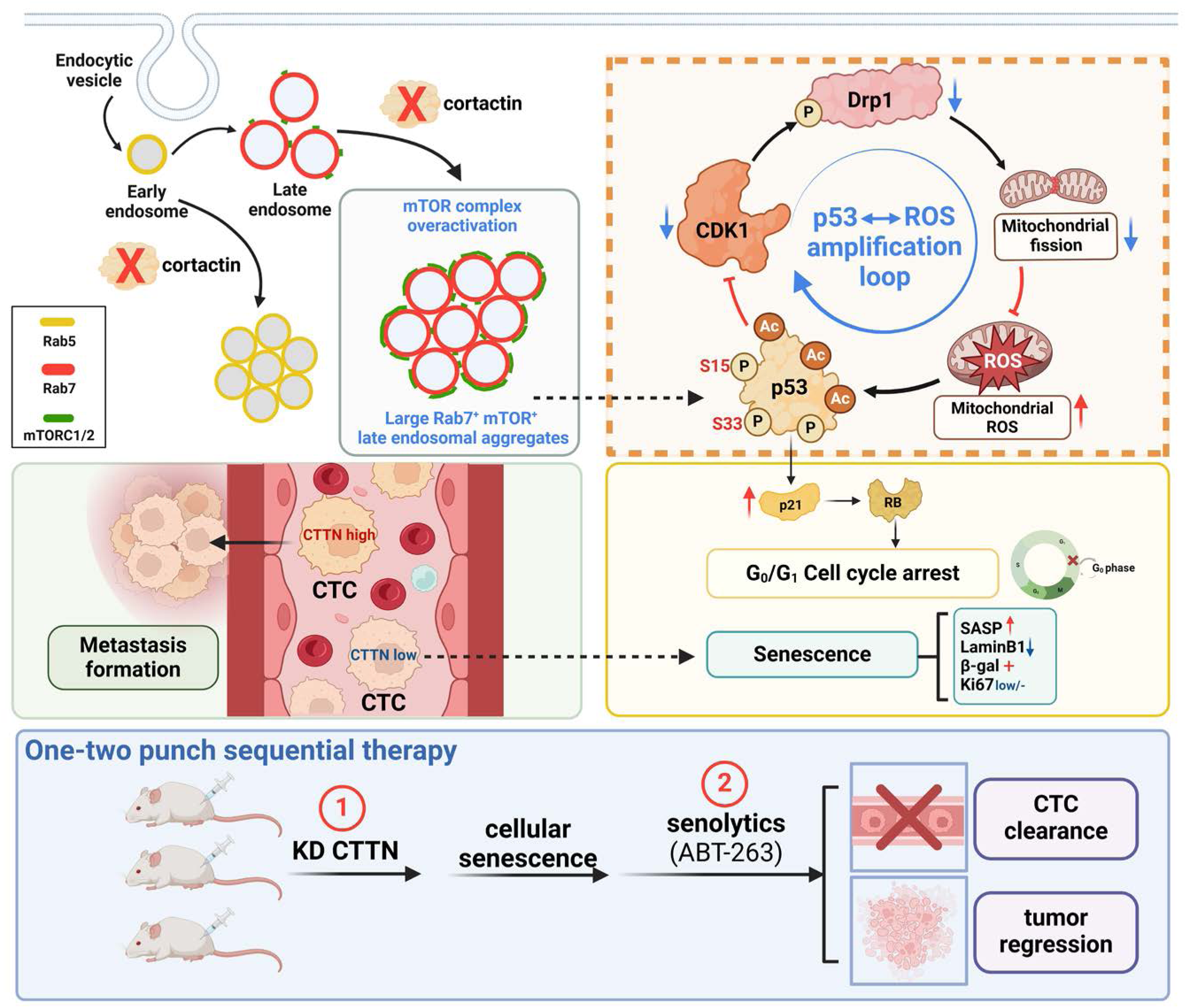
Working model illustrating the mechanisms of senescence regulation by cortactin and the “One-two punch” sequential therapy. Depletion of cortactin induced aberrantly enlarged Rab7-positive late endosomal aggregates within which mTOR was accumulated and overactivated. The activated mTOR further phosphorylated p53 on S15 and S33 sites. p53 and mitochondrial ROS signaling formed a “p53 ---| CDK1→p-DRP1 ---| mtROS→p53” amplification loop that enabled stable senescence induction and maintenance. A potential “One-two punch” sequential therapy was proposed for the treatment of metastasis by inducing cellular senescence of CTCs followed by the administration of senolytic drugs to eliminate these cells through the induction of apoptosis.

## Methods

### Cell culture

Mel-167 circulating tumor cell line established from melanoma patient blood ^39^ was cultured in RPMI-1640 medium (Gibico, 61870127) containing 20 ng/ml EGF(Gibico, PHG0313), 20 ng/ml bFGF (Gibico, 13256029), 1x B27 (Gibico, 17504044) at 4 % O2 and 5% CO2 atmosphere. A375 melanoma cell line and 293T was obtained from ATCC, cultured in Dulbecco’s Modified Eagle medium with high glucose (DMEM; Gibco, 11965118) containing 10% Fetal bovine serum (FBS; ExCell Bio, FSP500) at 5% CO2 atmosphere. All cell lines are routinely assayed for mycoplasma contamination.

### Antibodies, chemicals and sequences of primer, shRNA and siRNA

Information of antibodies, chemicals and sequences of primer, shRNA and siRNA can be found in **Table 1**-**3**.

**Table 1.**
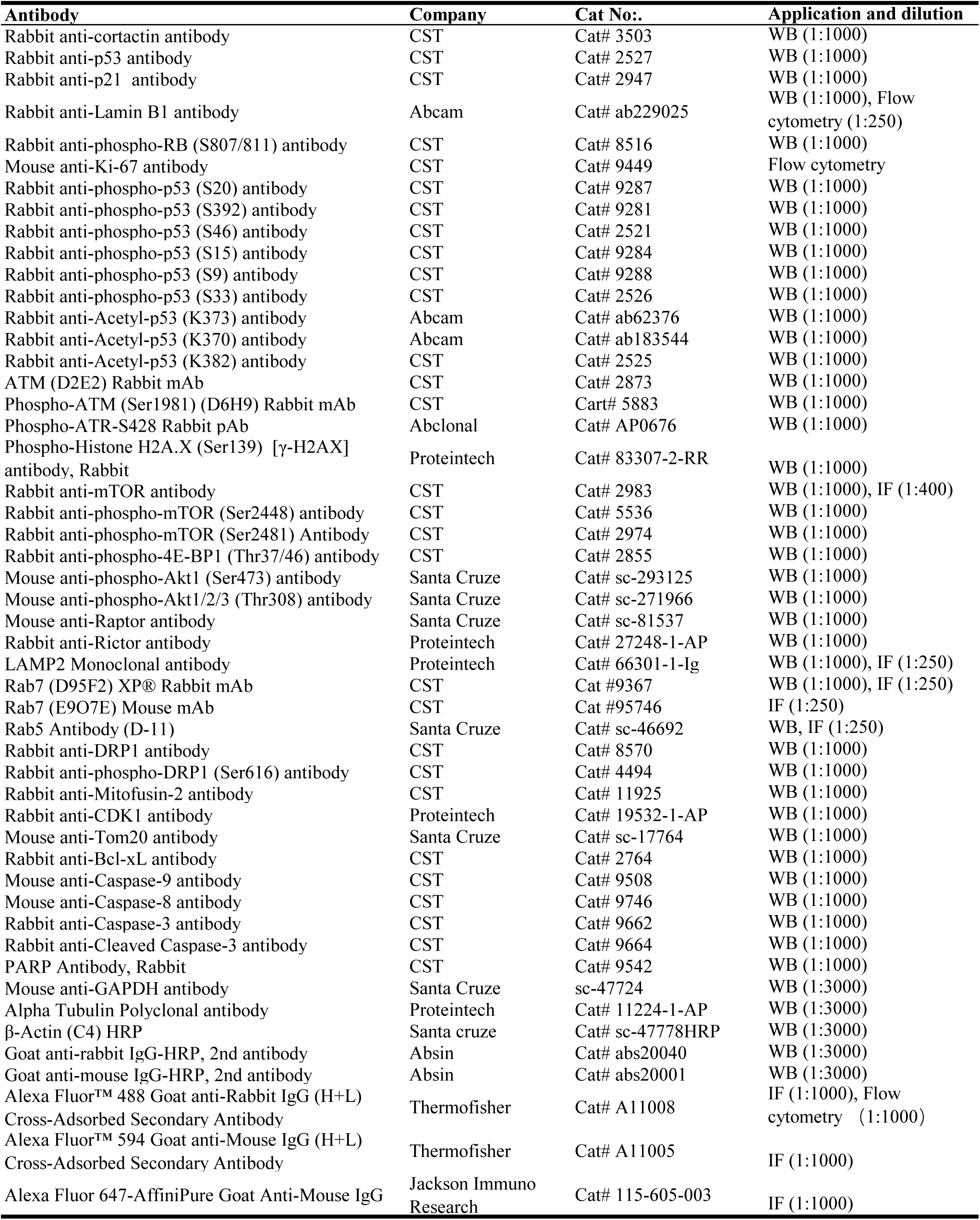
Antibody information.

**Table 2.**
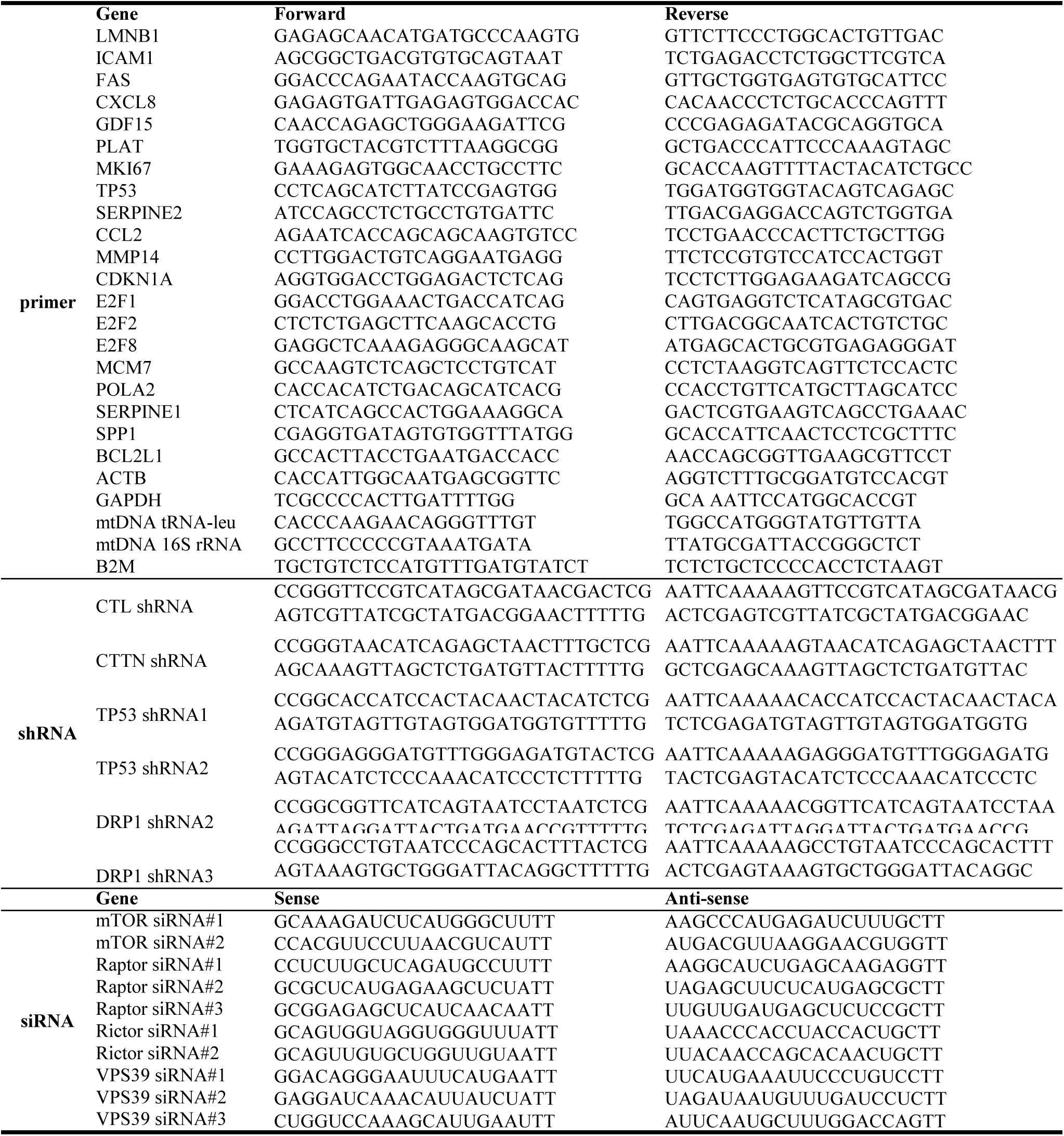
Primer, shRNA and siRNA sequences.

**Table 3.** Chemical list.

| Chemicals Name | Company | Catalog number | Function |
| --- | --- | --- | --- |
| Cycloheximide (CHX) | MCE | Cat# HY-12320 | Translation inhibitor |
| MG132 | MCE | Cat# HY-13259 | proteasome inhibitor |
| Dorsomorphin | MCE | Cat# HY-13418A | AMPK inhibitor |
| BI-D1870 | MCE | Cat# HY-10510 | RSK inhibitor |
| PNU112455A hydrochloride | MCE | Cat# HY-112468 | CKD5 inhibitor |
| CDK5-IN-3 | MCE | Cat# HY-145694 | CKD5 inhibitor |
| Doramapimod | Selleck | Cat# S1574 | p38MAPK inhibitor |
| SP600125 | Selleck | Cat# S1460 | JNK inhibitor |
| Enitociclib | MCE | Cat# HY-103019 | CDK9 inhibitor |
| AR-A014418 | Selleck | Cat# S7435 | GSK3β inhibitor |
| LDC4297 | Selleck | Cat# S7992 | CDK7 inhibitor |
| Cucurbitacin E | MCE | Cat# HY-N0417 | CDK1 inhibitor |
| Rapamycin | MCE | Cat# HY-10219 | Rapamycin (mTOR complex 1 inhibitor) |
| Torin 1 | MCE | Cat# HY-13003 | Torin1 (mTOR complex 1 and 2 inhibitor) |
| AZD-8055 | MCE | Cat# HY-10422 | AZD8005 (mTORC1/2 inhibitor) |
| Dactolisib | MCE | Cat# HY-50673 | Dactolisib (dual inhibitor of PI3K and mTORC1/2) |
| Wortmannin | MCE | Cat# HY-10197 | pan-PI3K inhibitor |
| Mdivi-1 | MCE | Cat# HY-15886 | DRP1 inhibitor |
| Mitoquinone mesylate (MitoQ) | MCE | Cat# HY-100116A | mitochondrial ROS scavenger |
| N-acetylcysteine (NAC) | Selleck | Cat# S1623 | ROS scavenger |
| Tempol | MCE | Cat# HY-100561 | ROS scavenger |
| Navitoclax (ABT-263) | MCE | Cat# HY-10087 | selective inhibitor of BCL2, BCL-XL and BCL-W |
| ABT-737 | MCE | Cat# HY-50907 | selective inhibitor of BCL2, BCL-XL and BCL-W |
| Doxycycline | MCE | Cat# HY-N0565 | to induce the expression of shRNA in teton-pLKO.1 vector |
| Propidium Iodide (PI) | Invitrogen | Cat# P3566 | dye used for cell cycle analysis |
| Hoechst 33342 | Invitrogen | Cat# H3570 | dye used for nucleus staining |
| MitoSOX Red | Molecular Probes | Cat# M36008 | dye used for mitochondrial ROS detection |
| JC-1 dye | Invitrogen | Cat# T3168 | dye used for mitochondrial membrane potential detection |

### Constructs and lentivirus preparation

p53 shRNA was cloned into pLKO.1-blast vector (Addgene, 8453), CTTN shRNA was cloned into Tet-pLKO-puro (Addgene, 21915) digested by AgeI and EcoRI restriction endonuclease. shRNAs were designed using the GPP Web Portal online tool (https://portals.broadinstitute.org/gpp/public/gene/search). CTTN CDS with HA tag at N-terminal was synthesized and cloned into the CD532A-1 vector between EcoRI, BamHI sites by Genewiz company.

To produce lentiviral supernatants, 7 µg lentiviral vectors (Tet-pLKO-puro, pLKO.1-blast or CD532A-1) were co-transfected with 7 µg psPAX2 and 2.4 µg VSV-G vectors to HEK293T cells, using polyethylenimine as transfection reagent. Sixty hours post transfection, lentiviral supernatants were filtered by 0.45 µm syringe filter (Thermo fisher 7232545). To infect cells, 0.3–0.75 ml lentivirus with 1.5 or 5 µg/ml polybrene for Mel-167 or A375, respectively, were added to tumor cells for 16 h in a well of 6-well plates. Then cells were subjected to 5-7 days’ puromycin or blasticidin selection to establish stable cell lines. For doxycycline inducible shRNA expression, 300 ng/ml doxycycline was used.

### Cell proliferation assay

For 2D culture or suspension culture, 5,000 A375 cells or 10,000 Mel-167 cells per well were seeded to 96-well plates (Corning, 3610) and cultured for 3 days. CellTiter-Glo® Luminescent Cell Viability Assay Kit (Promega, G7571) was used to quantify cell viability. 3D cultures were prepared as previously described ^87^. Briefly, 96-well plates was coated by growth factor-reduced Matrigel (Corning, 354230). 1500–3000 cells were seeded to each well in completed medium, containing 2% Matrigel. To quantify spheroid growth, CellTiter-Glo® 3D Cell Viability Assay Kit (Promega, G9682) was used to quantify cell viability according to the product manual.

### Cell cycle analysis

Cells were collected and wash by PBS for once, the cells were fixed by pre-cold 75% ethanol in −20 °C for overnight. After the fixation, cells were spin down and washed by PBS then cells were incubated with the Propidium Iodide (PI) staining buffer containing 0.1 % NP-40, 10 ug/ml RNase (Takara, 2158), 1:500 PI (Invitrogen, P3566) in PBS for 30 mins in dark. Flow cytometry (BD Accuri™ C6 FlowCytometer) was used to detect the PI signal using the PE-A channel. The cell cycle was analyzed by the Flowjo software.

### Immunoblotting

Cells were lysed in RIPA buffer (1% NP-40, 1% sodium deoxycholate, 0.1% SDS in PBS) with proteinase inhibitor cocktail (Roche, 04693132001) and phosphatase inhibitor cocktail (Roche, 04906837001) on ice for 30 mins. Cell extracts were pre-cleared by centrifugation at 13,000 × g at 4 °C for 10 min and protein concentration was determined by the BCA assay (Pierce, 23225) using a BioTek Synergy™ H1 Microplate Reader. Then lysates were resolved by SDS-PAGE gel electrophoresis and protein was transferred to PVDF membrane, followed by blocking, primary and secondary antibody incubation and signal were examined by using the enhanced chemiluminescence substrate (Pierce, 32106).

### Senescence β-Galactosidase staining assay

Senescence β-Galactosidase staining assay was performed according to the β-gal staining kit (Biosharp, BL133A). A375 treated with or without 300 ng/ml Doxycycline to knock down cortactin expression for 6 days, the cells were fixed by the Fixative Solution buffer then washed by PBS followed by β-Galactosidase staining solution incubation for overnight in dark in 37°C. Pictures were captured and β-gal staining positive cells were counted using image J software.

### RNA-seq

Mel-167 and A375 cells were treated with Doxycycline for 3 days or 4 days to knock down the expression of cortactin, then total RNA was extracted using the NucleoZOL reagent (Macherey-Nagel, #74040.200) according to the product manual. The RNA library construction and RNA-seq analysis was performed by the HaploX Genomics Center (Shenzhen, China). Experiment was performed with 2 biological replicates. Briefly, we (i)performed quality control of RNA-seq raw reads using FASTP ^88^, (ii)fastq files were mapped to the human reference genome hg38 and obtained read counts and TPM using STAR ^89^, (iii)identified differentially expressed genes in each condition through log_2_fold changes and adjusted p-values using DESeq2 ^90^.

### Pathway and gene enrichment analysis

The Enrichment Analysis (GSEA) approach was used with a p < 0.05 using the ‘clusterProfiler’ R package ^91^. The pathway and gene sets were downloaded from the public Molecular Signatures Database (MSigDB). The DESeq2package was used to identify significantly different pathways and gene sets across groups ^90^ .

### Proteomic sample preparation

The proteomics analysis was performed as previous description ^92,93^. The pellets were lysed with 500 μL lysis buffer (1% Sodium Deoxycholate (SDC, Sigma, D6750), 8M Urea, 100 mM Triethylammonium bicarbonate (TEAB, Sigma, T7408) and 1x protease inhibitor) and sonicated for 15 s on ice. Aliquots of 100 μg protein were diluted to 1 mg/mL with lysis buffer, 20 mM dithiothreitol (DTT, Sigma, D9779) were added and incubated for 30 min at 56°C, followed by alkylation with 40 mM iodoacetamide (IAA, Sigma, I1149) for 30 min at room temperature in dark, the proteins were digested with trypsin (Promega, V5111) at a ratio of enzyme: protein as 1: 50.

### LC-MS/MS analysis and data processing

The LC-MS/MS analysis was performed on an Orbitrap Eclipse Tribrid mass spectrometer (Thermo Fisher Scientific) coupled to an EASY-nLC 1200 system (Thermo Fisher Scientific). The protein quantitative information were obtained by label-free quantification with data-independent acquisition (DIA) model. Raw files were analyzed by DIA-NN version 1.8 ^94^ against the human database downloaded from SwissProt (20211231, 20375 entries). The false discovery rate (FDR) threshold was set as 1% and calculated by a decoy search. The fixed modification was carbamidomethylation of cysteines, and variable modifications were methionine oxidation and N-terminal acetylation, in addition, N-terminal methionine excision was also enabled. The missed cleavages parameter was set as 1 and accuracy for MS1 and MS2 was 5 ppm. Match between runs was enabled here.

### In vivo tumor growth and metastasis experiments

The animal studies were approved by the Institutional Animal Care and Use Committee of the Southern University of Science and Technology (IACUC NO. SUSTech-JY202402014) NCG mice (6-8 week old) were purchased from Gempharmatech Co., Ltd, Jiangsu province, China. To test tumor growth by subcutaneous tumor model, Mel-167 cells were collected and washed by PBS for twice. Then cells were resuspended in pre-cold 50% Matrigel in PBS on ice. 80x10^4^ cells was injected to the flank of NCG mice subcutaneously. Tumor size was examined every 3-5 days for the duration of the experiment. Tumor volumes were calculated using the formula V = (W^2^ × L)/2, where V is the tumor volume, W is the tumor width, and L is the tumor length.

To test tumor metastasis, cells expressing luciferase were collected and suspended in pre-cold PBS. 80x10^4^ cells in 0.1 ml were injected to mice through tail vein. Luciferase signal from metastatic cells were determined by IVIS® Spectrum In Vivo Imaging System (PerkinElmer). To knockdown cortactin expression in Mel-167 tumor, tet-on-pLKO-puro vector was used and drinking water containing 20 g/L sucrose and 1 g/L doxycycline was administered to induced the expression of cortactin shRNA. Drinking water was changed every 5-7 days.

### CTC isolation

1 ml cardiac blood from tumor metastasis mouse model at experiment end point was collected in EDTA anticoagulant tube, then CTC from mouse blood was isolated by the Celutriator TX1 microfluidic device (Shenzhen Genflow Technologies) ^95^ according to the manufacture’s manual. CTC number was counted based on the GFP using the fluorescence microscope, as Mel-167 cells express GFP.

### Immunohistochemistry (IHC)

Tissues were collected and fixed by formalin, and embedded by paraffin. IHC was performed as described in ^96^. 5 µm formalin-fixed paraffin-embedded breast tissue sections were prepared, dewaxed in xylene, rehydrated in serial downgraded alcohols (100%, 95%, 70%) and brought down to water. To retrieve antigen, sections were immersed in pre-heated 90–95 °C Envision Flex Target Retrieval Solution with high pH (DAKO, Denmark, K8004) and microwaved for further 15 min followed by naturally cool down. Endogenous peroxidase was blocked 3% H2O2 for 25 mins in dark at room temperature. Rabbit anti-p53 (1:100), Rabbit anti-phospho-RB (1:100), Rabbit anti-p21 antibody (1:100) was applied to the sections and incubated for overnight at 4 degree. REAL EnVision Detection System and DAB chromogenic substrate (DAKO, Denmark, K5007) were then incubated with sections for 30 and 2 min, respectively, at room temperature for signal detection. Counterstain was performed by hematoxylin staining. Image of the whole section was scanned by Pannoramic MIDI (3DHISTECH) using 20x objective. H-SCORE was calculated by Aipathwell software (Servicebio technology Co) to assess the expression level of p53, p21 and p-RB. H-SCORE=∑(pi×i)=(percentage of weak intensity ×1)+(percentage of moderate intensity ×2)+(percentage of strong intensity ×3).

### Immunocytochemistry (ICC)

Cells was fixed by 4% PFA at room temperature for 15 mins. Suspension cultured Mel-167 was spin down to glass slide after fixation using a Cytospin Cytocentrifuge (Thermo Fisher). Cells then was blocked and permeabilized by incubated with blocking and permeabilization buffer (5% normal goat serum, 0.1% TritonX-100 in PBS) for 40 mins, followed by primary antibody incubation for overnight at 4 degree. Fluorochrome conjugated secondary antibody (1:1000) was then applied to the slide or confocal plate for 1 hour. Then cells were stained with 1 μM Hoechst 33342 (Invitrogen, H3570) for 5 mins. LSM 980 or LSM 900 confocal fluorescence microscope (Zeiss) was used to capture the fluorescence signal.

### RT-qPCR

Total RNA was extracted using the NucleoZOL reagent (Macherey-Nagel, #74040.200) according to the product manual. RT was performed using EvoM-MLV Reverse transcription Kit (Accurate Biology, #AG11728) following the product manual. qPCR reaction was prepared by using the SYBR Green premix pro TaqHS Kit (Accurate Biology, #AG11718) and then qPCR was performed using a QuantStudio 12K Flex Real-Time PCR System (Applied Biosystems).

### Polysome isolation

Polysome was sedimented by sucrose cushion method modified from a previous established protocol ^97^. Briefly, cells were treated with 100 μg/ml CHX for 2 mins and collected and lysis in lysis buffer ( 20 mM Tris*Cl pH 7.4, 150 mM NaCl, 5 mM MgCl2, 1 mM DTT, 100 μg/ml cycloheximide, 1% Triton X-100, 25 U/ml Turbo DNase I, RNase inhibitor 1U/μL, 1x proteinase inhibitor and 1x phosphatase inhibitor) by triturating cells 15 times through a 27 gauge needle. Supernatant was saved by centrifugation for 10 minutes at 20,000 × g, 4°C. sample concentration was determined by Nanodrop using the RNA detection model. Then 200 μg samples were added to Ultra-Clear centrifuge tubes (Beckman Coulter, 344059) followed by adding 0.90 ml 1M sucrose cushion carefully at the very bottom of the tubes. Pelleting polysomes by centrifugation in a SW 41 Ti Swinging-Bucket Rotor (Beckman, 331362) at 39,000 rpm, 4°C for 15 hours using the Optima XE-100 Ultracentrifuge. The polysome pellet was saved and RNA was extracted by NucleoZOL reagent (Macherey-Nagel, #74040.200) and RNA was used for RT-qPCR to determine p53 mRNA level.

### Cycloheximide (CHX) chase assay

Cells were treated with 300 ng/ml doxycycline to induce CTTN shRNA expression to knock down CTTN mRNA for 3 days, followed by 8 nM Actinomycin D treatment for 4 hours, which was used to activate p53 as the p53 expression level of control cell examined by immunoblotting was limited. Then medium was refreshed with 50 μg/ml CHX treated for 0, 0.5, 0.75, 1, 2, 4, 7 hours to halt the protein synthesis. Cell lysates were prepared for immunoblotting to determine the protein level of p53 and GAPDH as loading control. Expression level was quantified by image J software.

### Lamin B1 expression analysis by flow cytometry

Mel-167 cells with or without cortactin and p53 knockdown were collected and fixed by 4% PFA at room temperature for 15 mins, followed by 90% methanol for permeabilization at −20 °C for overnight. Then cells were spined down, washed and incubated with Rabbit anti-Lamin B1 antibody (Abcam, ab229025) or Rabbit IgG isotype (CST, 3900) at room temperature for 1 hour followed by Alexa Fluor 488 Goat anti-Rabbit secondary antibody incubation for 40 mins. Cells were washed before flow cytometry analysis.

### Senescence score calculation (SENCAN)

SENCAN classifier online tool (https://rhpc.nki.nl/sites/senescence/classifier.php) was used to distinguish senescent and non-senescent cell, including cancer cells and fibroblasts, established by Jochems et al. ^98^. To calculate the senescence scores, gene expression matrix containing RNA sequencing read counts was uploaded to the SENCAN classifier.

### Mitochondria ultrastructure analysis by transmission electron

Transmission electron microscopy (TEM) analysis was performed by Wuhan Servicebio technology CO., LTD. Cells were collected and fixed in TEM fixative (Servicebio, G1102) at 4 °C overnight followed by fixation by 1% osmic acid in dark for 2 hours. Cell pellets were then dehydrated in an ascending series of ethanol, and embedded in EMbed 812 Resin (SPI, 90529-77-4). 60 nm ultrathin sections were prepared by an ultramicrotome (Leica, UC7). Ultrathin sections were stained with 2% uranium acetate in absolute alcohol for 8 mins and 2.6% lead citrate solution for 8 mins. Samples were then observed and images were captured by a transmission electron microscope (Hitachi, HT7700) at 60-80 kV. Mitochondria number per cell, mitochondrial cristae length, the ratio of cristae length to outer membrane length and cristae number per mitochondrion were calculated using the image J software.

### Mitochondrial DNA content analysis

Genomic DNA of cells were isolated using QIAamp DNA Mini Kit (Qiagen, 51304). qPCR reaction was prepared by using the SYBR Green premix pro TaqHS Kit (Accurate Biology, AG11718) and then qPCR was performed using a QuantStudio 12K Flex Real-Time PCR System (Applied Biosystems). Primers of mtDNA tRNA^leu^(UUR), mtDNA 16S rRNA and nuclear encoded B2M were designed. Relative mitochondrial DNA content was determined by the ratio of mtDNA tRNA^leu^(UUR) or mtDNA 16S rRNA to nuclear encoded B2M ^99^.

### Mitochondrial superoxide analysis (MitoSOX Red)

MitoSOX Red (Molecular Probes, M36008) was used to detected mitochondrial superoxide according to the product manual. Briefly, suspension cultured Mel-167 cells were spin down and washed by HBSS buffer (with Ca2+, Mg2+), then cells were incubated with 1μM MitoSOX Red for 30 mins in incubator. Cells was washed by HBSS buffer before flow cytometry analysis. Adhesion cultured A375 cells were washed by HBSS buffer (with Ca2+, Mg2+) and stained for 30 mins on plates, then trypsinized and washed by HBSS buffer followed by cytometry analysis.

### Mitochondria membrane potential analysis (JC-1)

JC-1 dye (Invitrogen, T3168) was used to detect the mitochondrial membrane potential. Briefly, cells were collected and washed by HBSS buffer (with Ca2^+^, Mg2^+^) and incubated with 2 μM JC-1 in incubator for 25 mins. Cells were washed by HBSS buffer and analyzed by flow cytometry using the FL1 and FL2 channel.

### Smart-seq2/3 data processing and analysis

For Fig 1a & b, public smart-seq2 datasets (GSE109761, GSE51827, GSE67980, GSE75367, GSE86978, PRJNA603782, PRJNA603789, PRJNA662599) were obtained from GEO. The Smart-seq2/3 data are aligned by STAR with genome reference hg38, then remove duplicate used picard, and indel realiment used GATK4. Smart-seq2/3 downstream analysis was based on the Seurat R package. Further quality control was applied to cells based on the following thresholds: 1) the number of expressed genes was greater than 300 and less than 5,000; 2) the cells contained less than 20% mitochondrial RNA. The DoubletFinder R package was applied to remove potential doublets. The filtered gene expression matrix for each sample was normalized and scaled by the “NormalizeData” and “ScaleData” functions in Seurat. The Harmony R package was used to adjust batch effects between different patients and integrate the gene expression matrices of all samples. Finally, we identified 56314 genes and assessed 291 CTCs. We performed principal component analysis (PCA) on the corrected expression matrix using highly variable genes (HVGs) identified by the “FindVariableGenes” function. Next, the “RunPCA” function was used to perform the PCA and the “FindNeighbors” function was used to construct a K-nearest-neighbor graph. The most representative principal components were used to determine different cell types with the “FindCluster” function. All CTCs were clustered unsupervised to obtain functional subclusters. To identify differentially expressed genes for each cell subtype, the “wilcoxauc” functions from the presto package were used with default parameters. The expression differences with *P* < 0.05 and log2(fold change, FC) > 0.3 were considered differentially expressed genes.

### scRNA-seq library preparation

Single-cell suspension was loaded onto microwell chip and processed by the Singleron Matrix® Single Cell Processing System. Barcoding Beads are subsequently collected from the microwell chip, followed by reverse transcription of the mRNA captured by the Barcoding Beads and to obtain cDNA, and PCR amplification. The amplified cDNA is then fragmented and ligated with sequencing adapters. The scRNA-seq libraries were constructed according to the protocol of the GEXSCOPE® Single Cell RNA Library Kits (Singleron) ^100^. Individual libraries were diluted to 4 nM, pooled, and sequenced on Illumina novaseq 6000 with 150 bp paired end reads.

### Data preprocessing and normalization

Raw reads from scRNA-seq were processed to generate gene expression matrixes using CeleScope (https://github.com/singleron-RD/CeleScope) v1.9.0 pipeline. Briefly, raw reads were first processed with CeleScope to remove low quality reads with Cutadapt v1.17 to trim poly-A tail and adapter sequences. Cell barcode and UMI were extracted. After that, we used STAR v2.6.1a ^89^ to map reads to the reference genome GRCh38 (ensembl version 92 annotation). UMI counts and gene counts of each cell were acquired with featureCounts v2.0.1 ^101^ software, and used to generate expression matrix files for subsequent analysis. Single-cell downstream analysis was based on the Seurat R package. Further quality control was applied to cells based on the following thresholds: 1) the number of expressed genes was greater than 150 and less than 6,000; 2) the cells contained less than 10% mitochondrial RNA. The DoubletFinder66 R package was applied to remove potential doublets. The filtered gene expression matrix for each sample was normalized and scaled by the “NormalizeData” and “ScaleData” functions in Seurat. Finally, we identified 36,601 genes and assessed 6,925 cells from CTC cell line MEL167.

### Computational scoring analysis of cycling CTCs *VS* non-cycling CTCs

CTCs with CTTN reads >0 were used for the analysis, CTCs were further reclassified into CTTN^high^ (Top 25%, n=147) and CTTN^low^ (Down 25%, n=147) groups based on the CTTN expression. S-phase and G2/M score were assessed by scoring cells for the expression of 43 S-phase-specific and 54 G2- or M-phase-specific genes ^102^. CTCs with summed scores >0 were considered as cycling CTCs while summed scores <=0 were considered as non-cycling CTCs ^44^ (refer to extended data Figure S2a-b).

### Patient survival rate analysis

Patient survival rate analysis were performed using the Kaplan–Meier method to estimate Overall survival (OS), FPI, or DSS, the Kaplan–Meier curves were compared using the log-rank test. Bulk RNA sequencing transcriptome data of patients with SKCM and the corresponding clinical data was downloaded from The Cancer Genome Atlas (TCGA) database (https://portal.gdc.cancer.gov/). The FPKM data were converted to TPM and merged with clinical data, after which patient-barcodes with no survival data or those with meaningless survival data were removed. Briefly, we (i) adopted a long-rank test for OS (Overall Survival) and DFS (Disease Free Survival) plot of TP53-WT melanoma and high metastatic patients (Stage IV) in TCGA. (ii)

Another big cohort of melanoma (SKCM, GSE65904) was also used for computing the association between the DMFS (Distant Metastasis-Free survival) and the expression of CTTN. (iii) The melanoma patients in TCGA were divided into high and low expression groups based on the median expression of CTTN KO-upregulate SASP associated genes. We then computed the connection between the overall survival rate and the expression groups. (iv) We also included two immunotherapy cohorts of melanoma (respectively accession number: ENA project PRJEB23709, dbGaP phs000452) and obtained the relative transcriptomic and clinical data from the online supplementary data appended to the published studies. By taking the median value of the CTTN KO-upregulate SASP associated genes in each patient, we divided the patients into high and low groups, and then applied a long-rank test for survival rate.

### Statistics and reproducibility

All data are from different biological replicates and independent repeats and are presented as mean ± s.e.m. Unpaired two-tailed Student’s t-tests were used for statistical analysis by GraphPad Prism 8 software. The correlation between the indicators was analyzed by Pearson’s coefficient. No statistical methods were employed to determine sample size a priori in the animal studies. Kaplan‒Meier analysis was used to calculate survival rate. *P*<0.05 was considered to be statistically significant.

## Data availability

The scRNA-seq and Bulk RNA-seq data of tumor cells generated in the present study have been deposited in the Genome Sequence Archive in BIG Data Center, Beijing Institute of Genomics, Chinese Academy of Sciences under BioProject: PRJCA031009 (publicly accessible at https://ngdc.cncb.ac.cn/gsa-human/s/Ig94kJcr, https://ngdc.cncb.ac.cn/gsa-human/s/9GW1Y2y8). The mass spectrometry proteomics data have been deposited to the ProteomeXchange Consortium via the PRIDE partner repository with the dataset identifier “PXD059177”. Reviewer access details Log in to the PRIDE website (https://www.ebi.ac.uk/pride/) using the following details: Project accession: PXD059177, Token: iSZibzThnoLB . The in-house dataset used in this study will be available upon reasonable request.

The public proteomic datasets of melanoma patients used in this study are available in (http://www.proteomexchange.org/) PXD: PXD001725, PXD001724, PXD009630, PXD017968, and PXD026086. The public transcriptomic data and clinical information from The Cancer Genome Atlas (TCGA) SKCM cohort were downloaded from the UCSC Xena data portal (https://xenabrowser.net). In addition, three public datasets can be obtained from (https://www.ncbi.nlm.nih.gov/) dbGap: phs000452, (https://www.ebi.ac.uk/ena) Project: PRJEB23709, and GEO: GSE65904. We also included the public smart-seq2 datasets (GSE109761 & GSE51827 & GSE67980 & GSE75367 & GSE86978 & PRJNA603782 & PRJNA603789 & PRJNA662599) from GEO. Any additional information required to reanalyze the data reported in this paper is available from the lead contact upon request.

## Acknowledgments

This research was funded by the National Natural Science Foundation of China (NSFC) fundings: 82173391, T2250610233 (to X.H.), 32200643 (to J.H.); Guangdong provincial funding awards: 2021QN02Y112, 2023A1515010287 (to X.H.); Shenzhen Science and Technology Innovation Commission fundings: RCBS20221008093246069 (to J.H.), Guangdong Basic and Applied Basic Research Foundation: No. 2023B1515120036 (to L.D.); Science and Technology Project of Shenzhen (No. JCYJ20220530154407017 to H.Y., JCYJ20240813155824032 to H.Y.); Science and Technology special fund of Hainan Province (No. ZDYF2024SHFZ045 to H.Y. and P. Z.). We thank members of the Hong’s lab for their helpful comments on this manuscript. We thank the Life Science Research Facility of Core Research Facilities, Laboratory Animal Center and the Center for Computational Science and Engineering of SUSTech for providing strong support.

## Author Contributions

J.H., H.Y. and X.H. conceived, designed, and supervised the study. B.Z., J.C., G.H., H.Z., F.Y., K.L., S.Z., X.L., J.L., H.H., L.W., J.Z., L.D., Q.C., H.L., J.H., H.Y. and X.H. carried out experiment, data analysis and wrote the original manuscript. All authors reviewed and revised the manuscript. All authors have read and agreed to the published version of the manuscript.

## Competing interests

The authors declare no competing interests.

---

**Extended Data Fig. 1.**
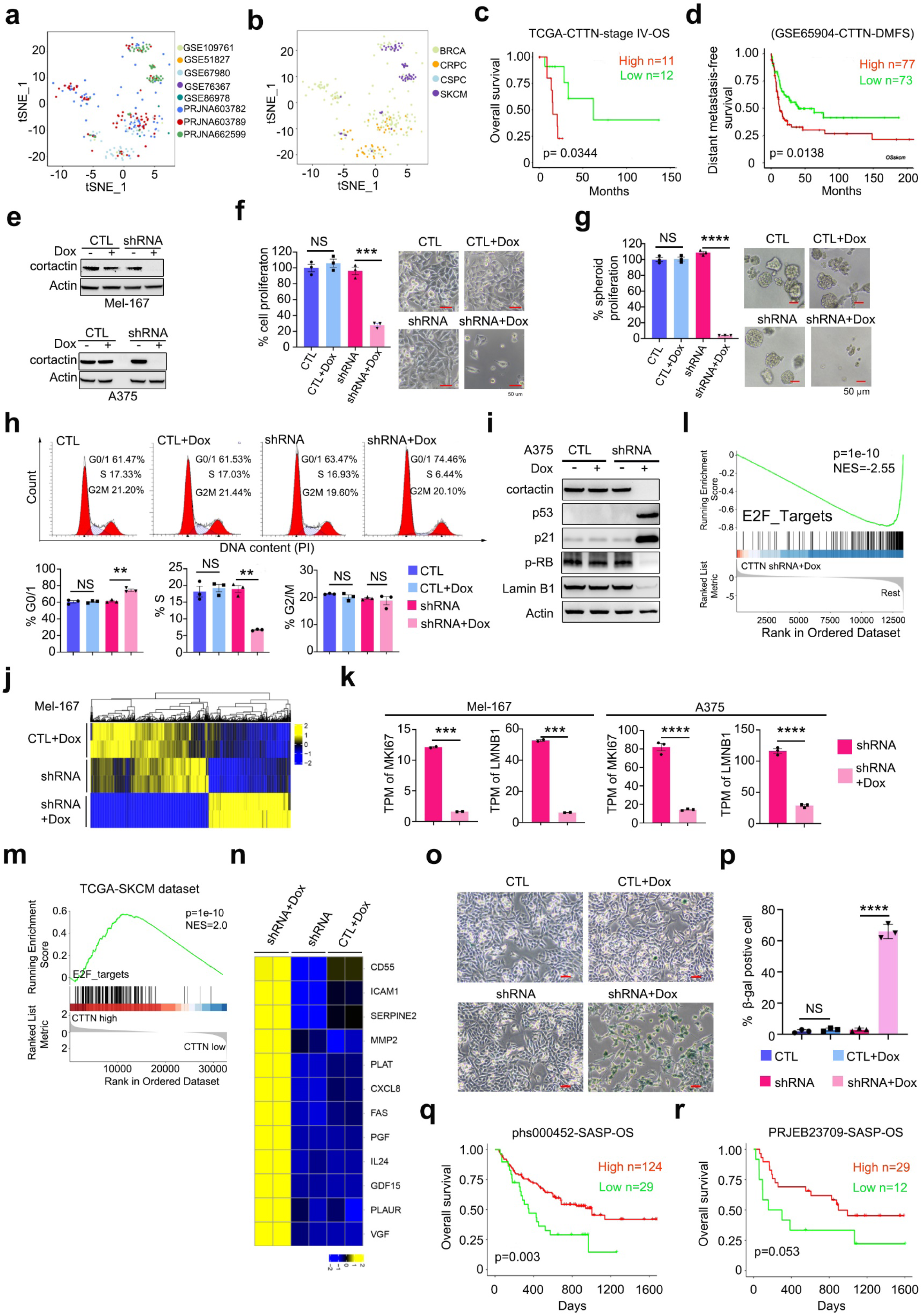
Cortactin depletion induces senescence in CTCs and metastatic melanoma A375 cells. **a**, t-SNE map of 291 CTCs analyzed by scRNA-seq across all datasets. Clusters were annotated by the dataset’s origin. Datasets were indicated. **b,** t-SNE map of 291 CTCs analyzed by scRNA-seq across all datasets. Clusters were annotated by cancer types. BRCA, breast cancer; CRPC, castration-resistant prostate cancer ; CSPC, castrate-sensitive prostate cancer; SKCM, skin cutaneous melanoma. **c,** Overall survival of patients with stage IV SKCM, analyzed by using the TCGA-SKCM data set. n=11 and 12 for CTTN high and low, respectively. **d,** Distant metastasis-free survival (DMFS) of patients with SKCM, analyzed by using the GSE65904 data set. n=77 and 73 for CTTN high and low, respectively. **e,** Immunoblotting detection of the CTTN KD efficiency by doxycycline-inducible shRNA system in Mel-167 (upper panel) and A375 (lower panel). 300 ng/ml doxycycline treatment for 3 days. **f,g**, Quantification of cell proliferation of A375 cell line cultured in conventional 2D (**f**) and 3D model (**g**). Pictures showed the representative morphology of cells with or without CTTN KD. n=3 independent experiments. Scale bars, 50 μm. **h**, Cell cycle analysis of A375 by Propidium Iodide (PI) staining followed by flow cytometry. Bar graphs showed the quantification of the percentage of G0/G1, S and G2M phase. n=3 independent experiments. **i**, Immunoblotting detection protein level of Cortactin and senescence marker, p53, p21, phosphorylated-RB (p-RB) and Lamin B1 of A375. Doxycycline treatment for 3 days. **j**, Heatmap portrayed the transcriptomic changes in Mel-167 CTC with or without CTTN KD for 3 days. n=2 biological repeats. **k,** mRNA level of MKI67 (Ki-67) and LMNB1 (Lamin B1) of Mel-167 and A375 with or without CTTN KD, detected by RNA-seq. n=2 (Mel-167) or 3 (A375) biological replicates. TPM, Transcripts Per Million. **l**, Gene set enrichment analysis showed that the downregulation of E2F_targets in CTTN KD-Mel-167 cells. **m,** Picture showed the GESA of E2F targets using TCGA-SKCM dataset comparing CTTN mRNA high and low expressed samples. **n**, Heatmap showed significantly upregulated SASP genes of Mel-167 upon CTTN KD by doxycycline induction for 3 days. Cutoff, *P* values <0.05, log2FC >0.58. n=2 biological repeats. **o,** Representative picture showed the SA-β-gal staining of A375 cells with or without CTTN KD for 6 days. **p,** quantification of SA-β-gal stained positive cell of (**n**). n=3 independent experiments **q,** Kaplan–Meier overall survival curve, stratificated by high and low score of CTTN KD-induced SASP signature. melanoma patient samples from dataset phs000452. **r,** Kaplan–Meier overall survival curve, stratificated by high and low score of CTTN KD-induced SASP signature. melanoma patient samples from dataset PRJEB23709. Dox, doxycycline; CTL, doxycycline inducible control shRNA; shRNA, doxycycline inducible CTTN-targeting shRNA. Data are represented as mean ± SEM in (**f**-**h**), (**k**) and (**p**). The statistical significance was calculated by two-sided Student’s t test in (**f**-**h**), (**k**) and (**p**). \*\**P*<0.01; \*\*\**P*<0.001; **** *P*<0.0001; NS, not significant. *P* values of (**c,d**), (**q,r**) were calculated by log rank test.

**Extended Data Fig. 2.**
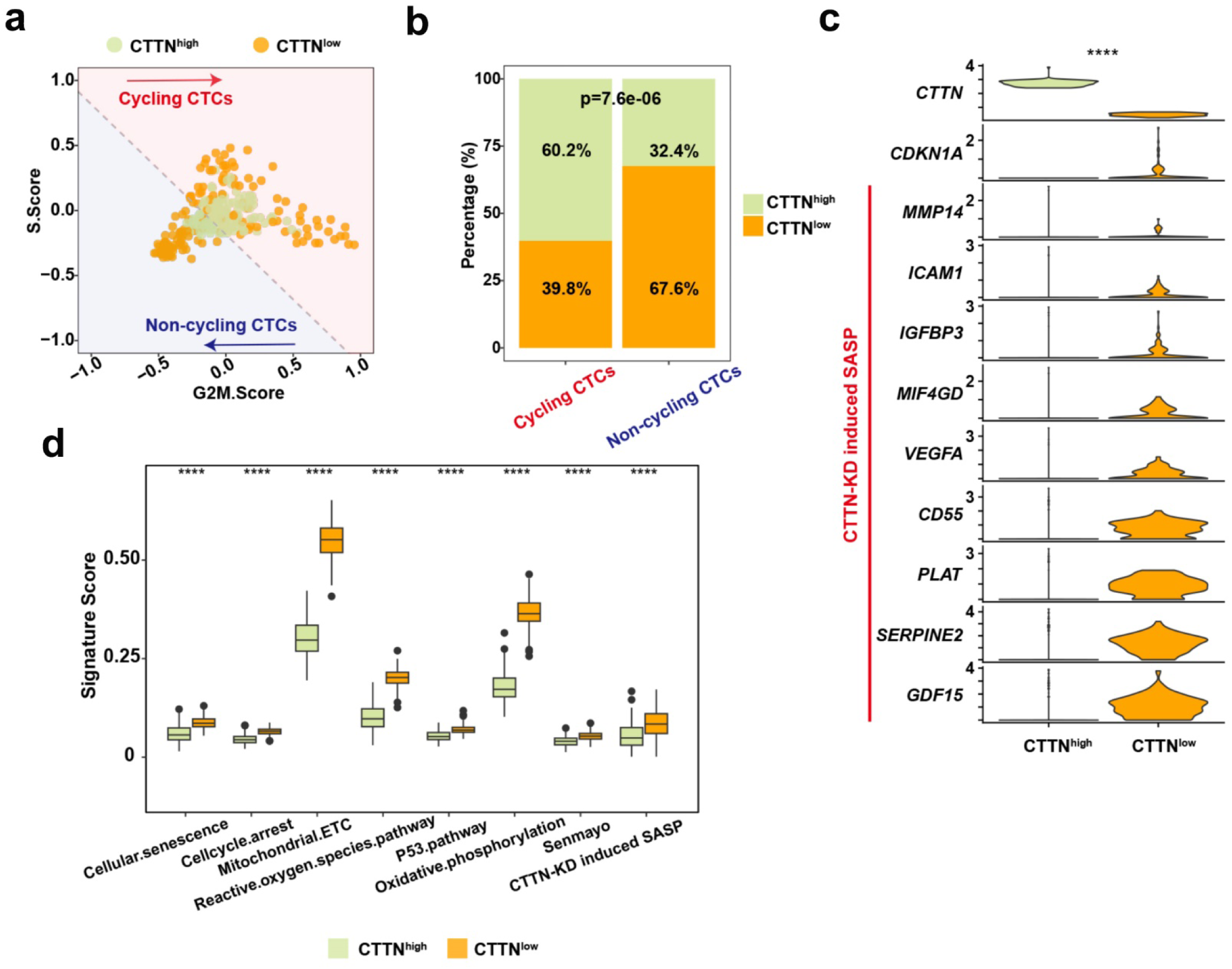
Single-cell RNA-seq of early CTC cultures reveals heterogenous expression of CTTN in single CTCs that are inversely correlated with the senescence state. **a**, Scatter plot of S- and G2M-phase gene expression scores for individual cells extracted on Mel-167 cell line and colored by CTTN-high and CTTN-low subpopulations. Dotted lines indicate thresholds of cells with score sums >0. **b**, Bar plot showing the proportion of cell cycle status (cycling, non-cycling) in CTTN-high and CTTN-low cells in a. **c**, Violin plot showing the senescence-associated genes expression in CTTN-high and CTTN-low cells. *P* value in (**c**) was calculated using a two-sided paired Wilcoxon signed-rank test. **d**, Boxplot showing the expression level of senescence-associated signature scores in CTTN-high versus CTTN-low cells. Data are presented as the mean ± SD. **** *p* < 0.0001, two-tailed Student’s t-test.

**Extended Data Fig. 3.**
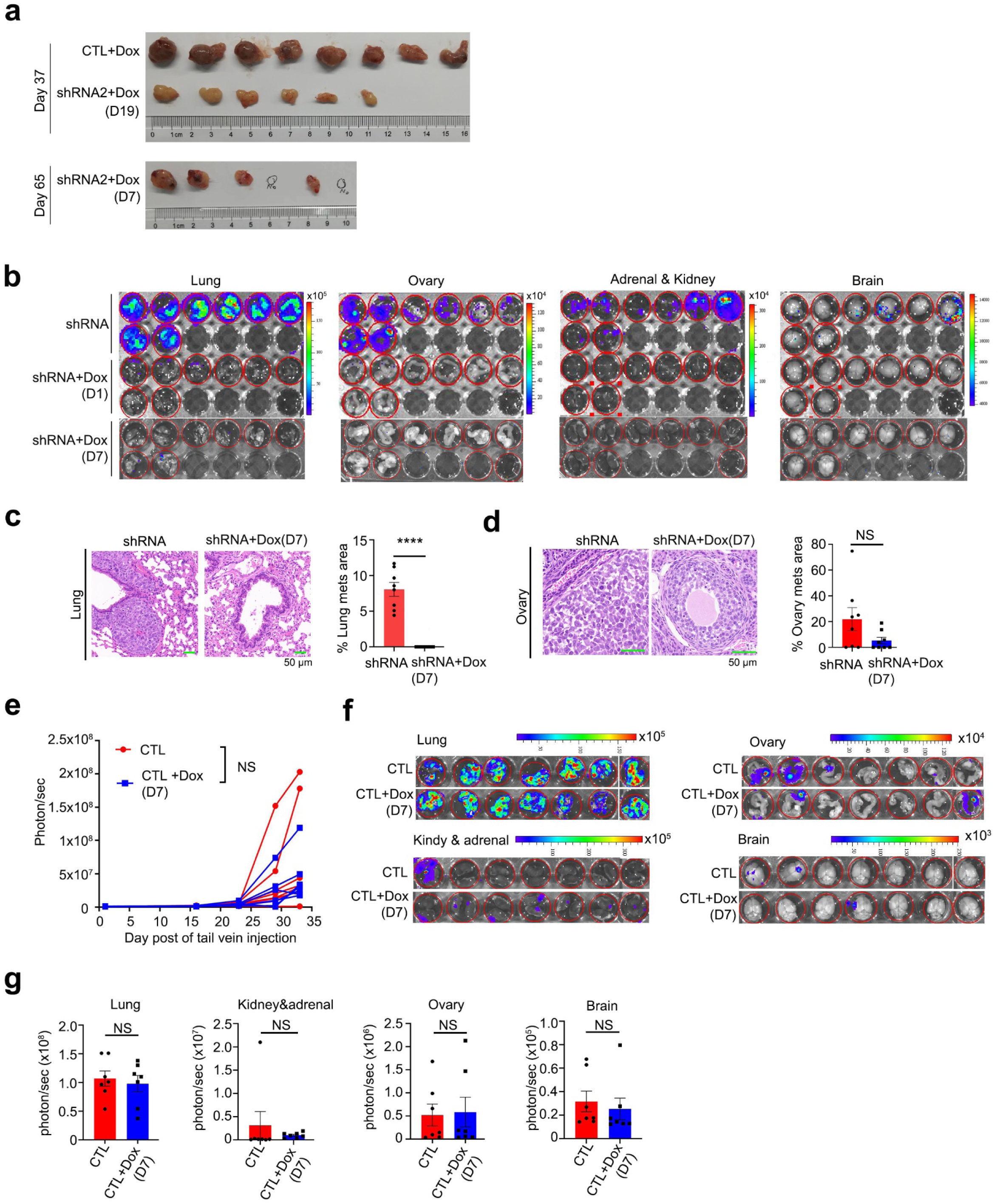
Cortactin depletion inhibits melanoma tumor growth and metastasis. **a**, Picture showed the tumors dissected from Mel-167 CTC s.c. xenograft model with or without doxycycline (Dox) to knock down CTTN expression. 2 tumors treated with Dox at day 7 were disappeared at day 65. n (CTL +Dox) =8 mice; n (other 2 group) =6 mice. **b**, Ex vivo IVIS imaging of lung, ovary, adrenal and kidney and brain dissected from metastasis mice model of Mel-167 with or without CTTN KD by Dox treatment. n=8 mice. **c, d**, H&E staining of lung (**c**) and ovary (**d**) dissected from the metastasis mice model of Mel-167 with or without CTTN KD. n=8 mice. **e**, Metastatic tumor growth curve of Mel-167-teton CTL shRNA with or without Dox treatment, luciferase signal was detected by IVIS. n=7 mice. **f, g**, Picture showed the ex vivo IVIS signal of lung and ovary metastatic sites of Mel-167-teton CTL shRNA with or without Dox treatment (**f**). Bar graph (**g**) showed the quantification of luminescence signal of metastatic tumor cells. n=7 organs. Dox, doxycycline. D7, D19, doxycycline administrated on day 7 and day 19, respectively. Data are represented as mean ± SEM in (**c, d**) and (**g**). The statistical significance was calculated by two-sided Student’s t test in (**c-e**) and (**g**). **** *P*<0.0001; NS, not significant.

**Extended Data Fig. 4.**
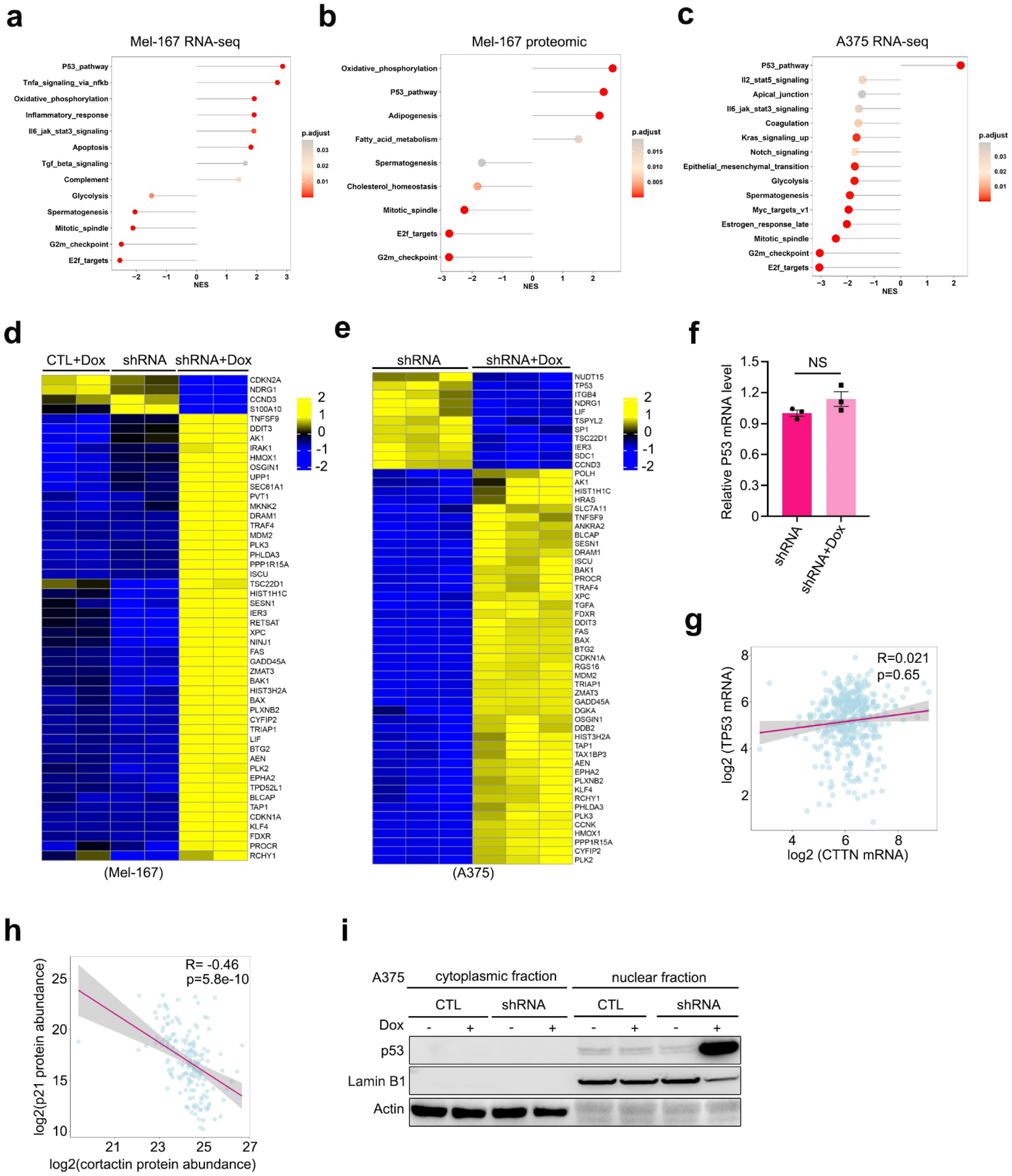
Transcriptomic and proteomic changes of melanoma cells upon cortactin depletion. **a-c**, The lollipop plot showed Hallmark50 pathways enrichment between control and CTTN KD in Mel-167 at transcriptomic level (A) and proteomic level (B) and A375 at transcriptomic level (**c**). NES, normalized enrichment score. Doxycycline treatment for 4 days. **d, e,** Heat map showed the significant mRNA changes of p53 pathway genes of Mel-167 (**d**) and A375 (**e**) treated by doxycycline for 3 and 4 days respectively, to knock down CTTN. Cutoff, *P* values <0.05, log2FC >0.58 and log2FC<-0.58. **f**, Relative p53 mRNA level of Mel-167 determined by RT-qPCR. n=3 independent experiments. **g**, The mRNA expression correlation between TP53 and CTTN determined by Pearson’s correlation analysis, using the TCGA-SKCM dataset. **h**, The protein expression correlation between p21 and cortactin determined by Pearson’s correlation analysis, using the human melanoma proteome dataset from Lazaro et al cohort (PMID: 34323403). **i**, Immunoblotting of p53 from cytoplasmic fraction and nuclear fraction of A375 with or without CTTN depletion. *P* value of (**g, h**) was calculated by Pearson correlation test.

**Extended Data Fig. 5.**
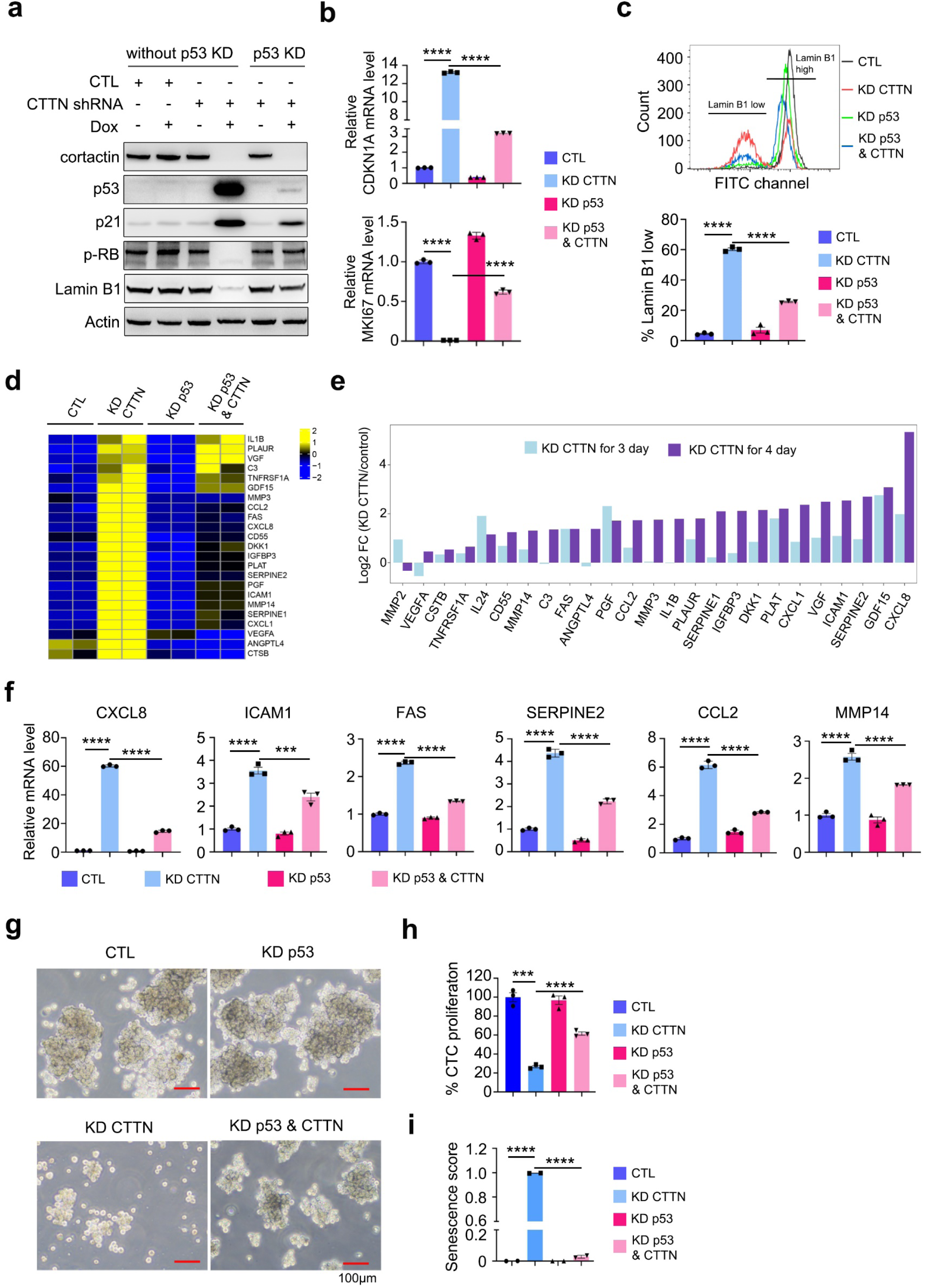
KD CTTN-induced cellular senescence and growth suppression is p53-dependent. **a**, Immunoblotting of senescence markers protein level of Mel-167 with or without CTTN and or p53 knockdown. CTTN shRNA was cloned into teton-pLKO.1_puro vector, p53 shRNA was cloned into pLKO.1_Blast vector. **b**, CDKN1A (p21) and MKI67 (Ki-67) mRNA level of Mel-167 determined by qPCR. n=3 independent experiments. **c**, Flow cytometry analysis of Lamin B1 protein level of Mel-167 with or without CTTN and p53 depletion. Bar graph showed the quantification of the percentage of Lamin B1 low Mel-167 cell. n=3 independent experiments. **d**, Heatmap showed the differential expression SASP genes of Mel-167 with or without CTTN and or p53 knockdown. Cells were treated by doxycycline for 4 days to KD CTTN. Cutoff, *P* value <0.05, log2FC >0.58 and log2FC <-0.58. **e,** Bar graph displayed the log2 Fold change of SASP genes showed on Extended Data Fig. 1o (3 days doxycycline induction) and Extended Data Fig. 5d (4 days doxycycline induction). **f**, RT-qPCR detection the mRNA level of SASP genes, CXCL8, ICAM1, FAS, SERPINE2, CCL2 and MMP14 in Mel-167. n=3 independent experiments. **g**, Representative pictures showed the Mel-167 cells cultured in suspension condition with or without CTTN and or p53 depletion **h**, Quantification of Mel-167 CTC proliferation of (**g**), n=3 independent experiments. **i**, Senescence score of Mel-167 calculated by the SENCAN method using RNA-seq data (same RNA-seq date of Extended Data Fig. 5d). n=2 biological repeats. Data are represented as mean ± SEM in (**b,c**), (**f**) and (**h,i**). The statistical significance was calculated by two-sided Student’s t test in (**b,c**), (**f**) and (**h,i**). \*\*\**P*<0.001; **** *P*<0.0001.

**Extended Data Fig. 6.**
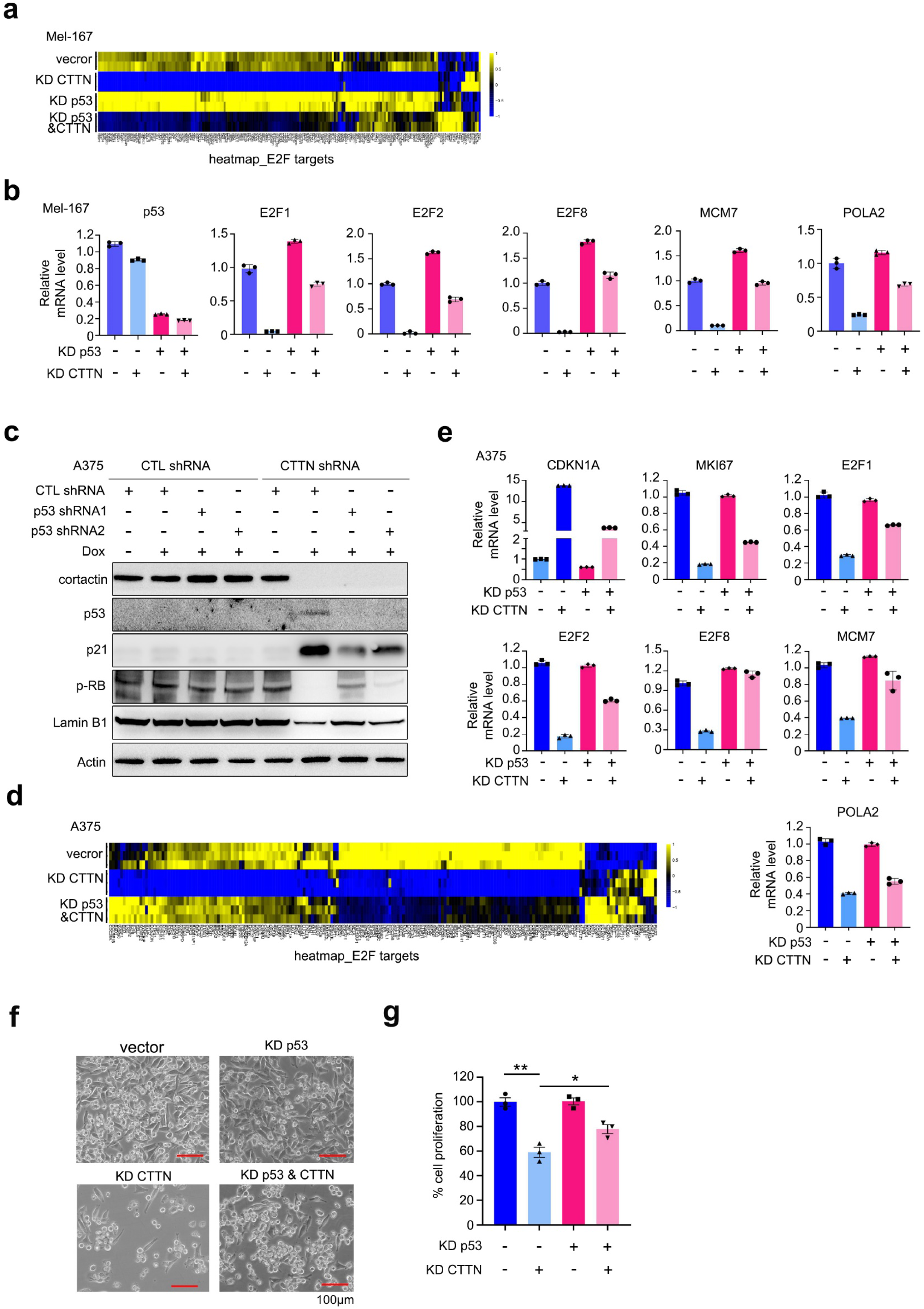
KD CTTN induces p53-dependent changes in E2F targets and the senescence induction in A375 cells. **a**, Heatmap showed the differential expression gene of E2F targets of Mel-167 with or without CTTN and p53 knockdown. Cells were treated by doxycycline for 4 days to KD CTTN. n=2 biological repeats. **b**, Relative mRNA level of p53, E2F1, E2F2, E2F8 and E2F targets MCM7 and POLA2 of Mel-167 with or without CTTN and p53 knockdown. n=3 technical repeats. **c**, Immunoblotting of senescence markers level in A375 with or without CTTN and or p53 knockdown. Doxycycline for 4 days to KD CTTN. **d**, Heatmap showed the differential expression gene of E2F targets of A375 with or without CTTN and p53 knockdown. Doxycycline for 4 days to KD CTTN. n=2 biological repeats. **e**, Relative mRNA of CDKN1A, MKI67, E2F1, E2F2, E2F8 and E2F targets MCM7 and POLA2 of A375 with or without CTTN and p53 knockdown. n=3 technical repeats. **f**, Representative pictures showed A375 cells cultured in 2D with or without CTTN and or p53 depletion. **g**, Quantification of A375 proliferation of (**f**), n=3 independent experiments. Data are represented as mean ± SEM in (**b**), (**e**) and (**g**). The statistical significance was calculated by two-sided Student’s t test in (**g**). * *P*<0.05; ** *P*<0.01.

**Extended Data Fig. 7.**
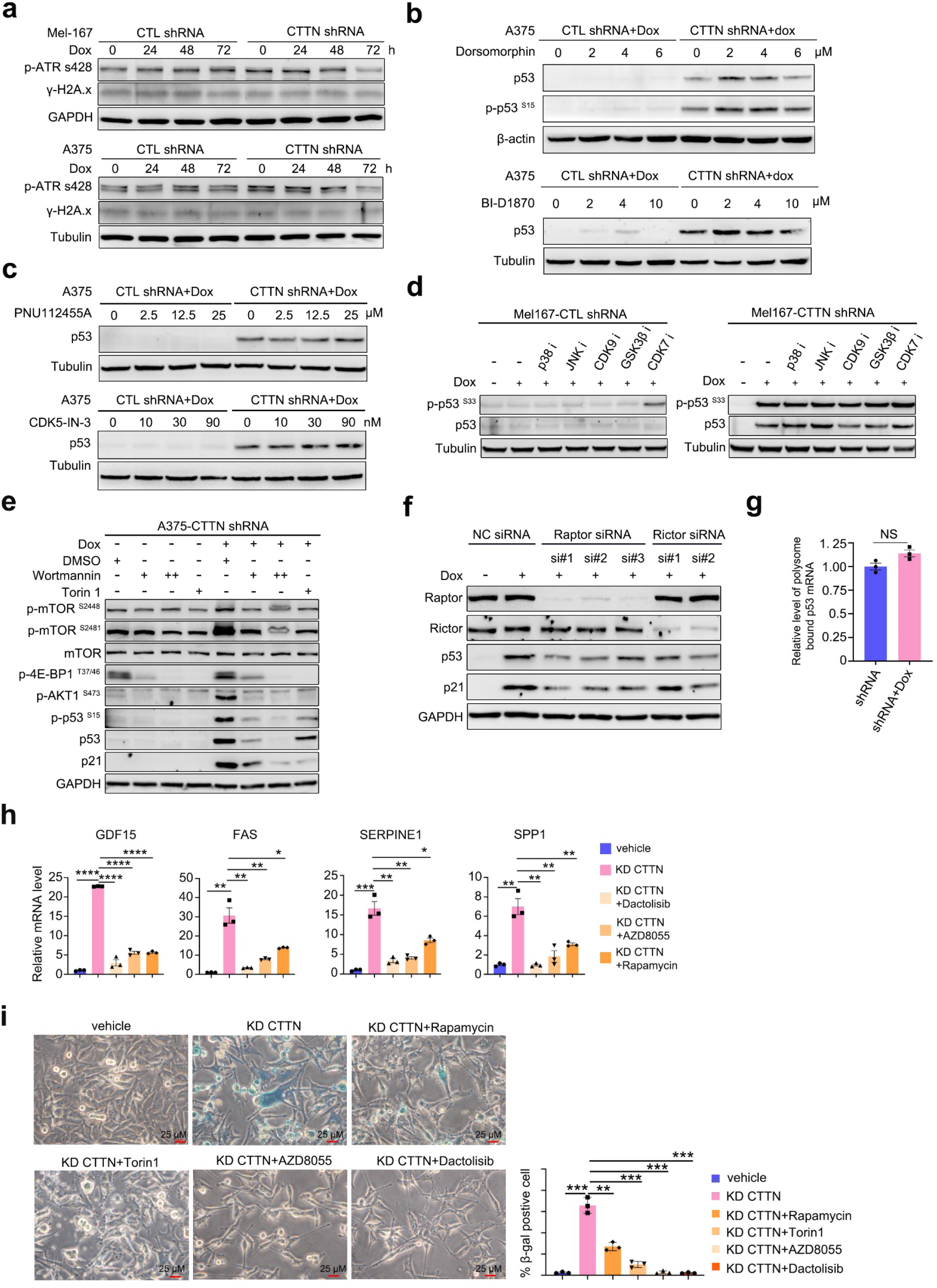
Candidate-based kinase screening identifies mTOR as the major upstream regulator of p53 pathway activation in CTTN-KD cells. **a,** Protein level of phosphorylated ATR S428 and γ-H2A.x upon CTTN depletion in Mel-167 (upper panel) and A375 cells (lower panel). **b,** Protein level of phosphorylated p53 S15 and p53 upon CTTN depletion in A375 cells in the present of AMPK inhibitor Dorsomorphin (upper panel) or RSK inhibitor BI-D1870 (lower panel) treatment for 48h. **c,** Protein level of p53 upon CTTN depletion in A375 cells in the present of CDK5 inhibitor PNU112455A (upper panel) or CDK-IN-3 (lower panel) treatment for 48h. **d,** Protein level of phosphorylated p53 S33 and p53 upon CTTN depletion in Mel-167 in the present of p38MAPK inhibitor (25 μM Doramapimod), JNK inhibitor (10 μM SP600125), CDK9 inhibitor (30 nM Enitociclib), GSK3β inhibitor (2.5 μM AR-A0144), CDK7 inhibitor (1 μM LDC4297) for 48h. **e,**Immunoblotting detection of mTOR, p-mTOR S2448, S2481, p-4E-BP1 T37/46, p-AKT1 S473, p-p53 S15, p53 and p21 of A375-teton CTTN shRNA treated with or without 48h doxycycline and 200 nM Wortmannin or 250 nM Torin1 for the last 4 hours. **f,** Immunoblotting detection of Raptor, Rictor, p53, p21 upon Raptor or Rictor knockdown by siRNA for 48h in A375-teton CTTN shRNA cells with or without doxycycline treatment to KD CTTN. **g,** RT-qPCR determination of p53 mRNA level followed by extraction RNA from isolated polysome from Mel-167 with or without CTTN depletion. n=3 independent experiments. **h,** RT-qPCR quantification of SASP related gene expression level of A375 with or without CTTN KD and or mTOR inhibitor treatment. n=3 independent treatment. **i,** Representative picture showed the SA-β-gal staining of A375 cells with or without CTTN KD for 5 days and mTOR inhibitors treatment for 3 days. Bar graph showing the quantification of the percentage of SA-β-gal positive cells. n=3 independent experiments. Data are represented as mean ± SEM in (**g**) and (**h,i**). The statistical significance was calculated by two-sided Student’s t test in (**g**) and (**h,i**). * *P*<0.05; ** *P*<0.01; *** *P*<0.001; **** *P*<0.0001; NS, not significant.

**Extended Data Fig. 8.**
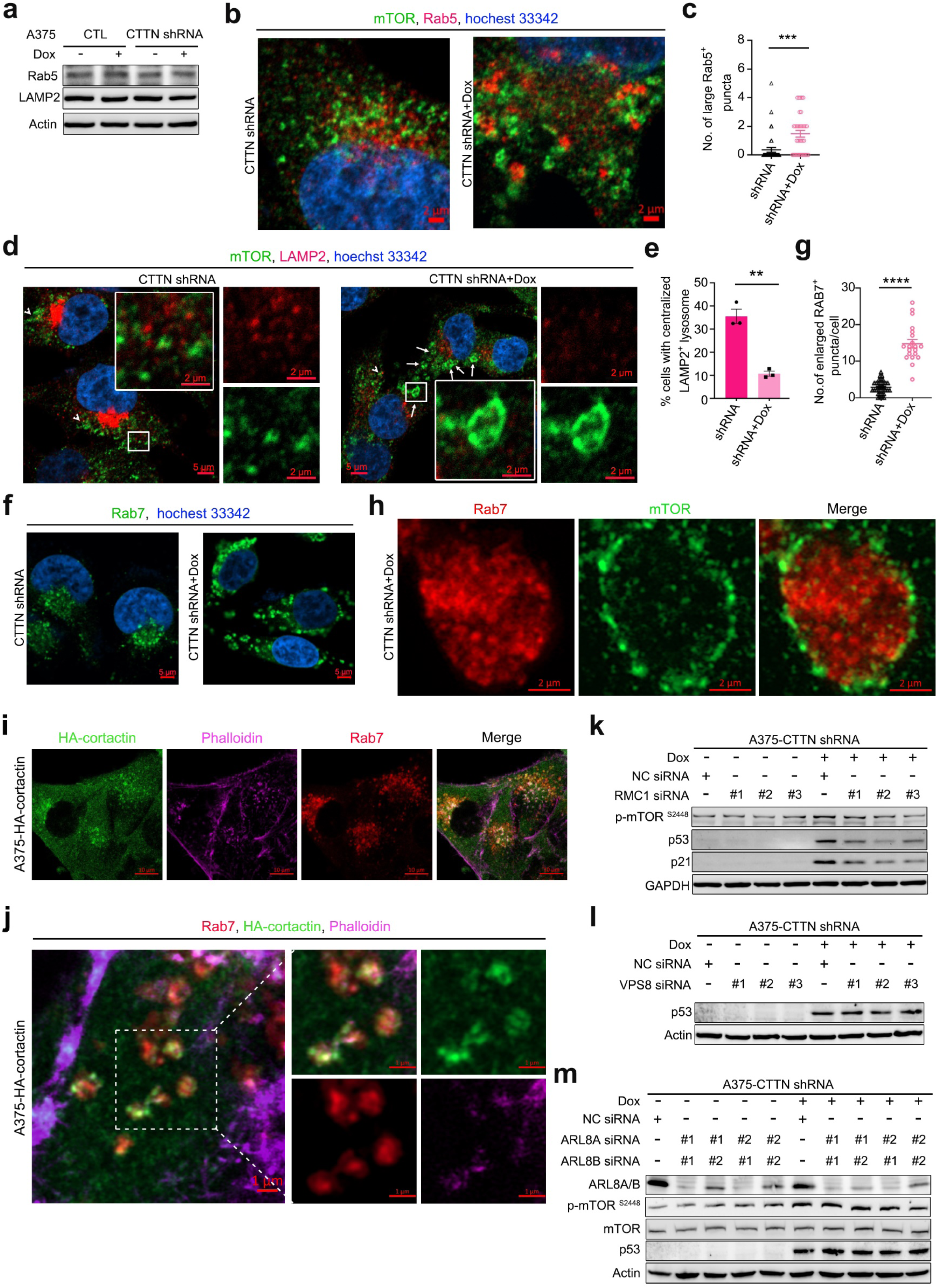
KD CTTN leads to aberrant Rab7-positive late endosomal aggregates with mTOR overactivation. **a,** LAMP2, Rab5 protein level of A375 with or without CTTN KD, determined by immunoblotting. **b,c,** Confocal imaging of mTOR, Rab5 of A375 cells with or without CTTN KD for 2 days (**b**). Scatter plot showed the quantification of large Rab5 positive puncta per cell (**c**). Puncta> 1.5 μm were counted. n=39 and 35 cells, with and without CTTN KD, respectively. Scale bar, 2 μm. **d,** Confocal imaging of mTOR and lysosome marker LAMP2 of A375 cells with or without CTTN KD for 2 days. Nuclei were stained by Hoechst 33342. Arrows indicate the large vesicle-like mTOR structures. Arrow heads indicate puncta with colocalization of mTOR and LAMP2. Scale bar, 5 μm and 2 μm. **e**, Bar graph showed the percentage of A375 cells with centralized LAMP2 positive lysosome distribution upon CTTN KD for 2 days. n=3 independent experiments. **f,g,** Confocal imaging of Rab7 of A375 cells with or without CTTN KD for 2 days (**f**). Scatter plot showed the number of large Rab7 positive puncta (>1.5 μm) per cell (**g**). n= 35 and 20 cells with and without CTTN KD, respectively. Scale bar, 5 μm. **h,** Immunofluorescence of mTOR and Rab7 of A375 cells with or without CTTN KD for 2 days. Super-Resolution Image was captured by using the Airyscan 2 model of ZEISS LSM 900. Scale bar, 2 μm. **i**, Immunofluorescence of Rab7, HA-cortactin (anti-HA tag) and F-actin (phalloidin) of A375 cells expressing HA-cortactin. Images were captured by confocal model. Scale bar, 10 μm. **j,** Immunofluorescence of Rab7, HA-cortactin (anti-HA tag) and F-actin (phalloidin) of A375 cells expressing HA-cortactin. Super-Resolution Images were captured by using the Airyscan 2 model of ZEISS LSM 900. Scale bar, 1 μm. **k,** Immunoblotting detection of protein levels of p-mTOR S2448, p53 and p21 in A375 cells with or without CTTN and/or RMC1 knockdown for 48h. **l,** Immunoblotting detection of protein levels of p53 in A375 cells with or without CTTN and/or VPS8 knockdown for 48h. **m,** Immunoblotting detection of protein levels of ARL8A/B, p-mTOR S2448, mTOR and p53 in A375 cells with or without CTTN and/or ARL8A/B knockdown for 48h. Data are represented as mean ± SEM in (**c**), (**e**) and (**g**). The statistical significance was calculated by two-sided Student’s t test in (**c**), (**e**) and (**g**). \*\**P*<0.01; \*\*\**P*<0.001,**** *P*<0.0001.

**Extended Data Fig. 9.**
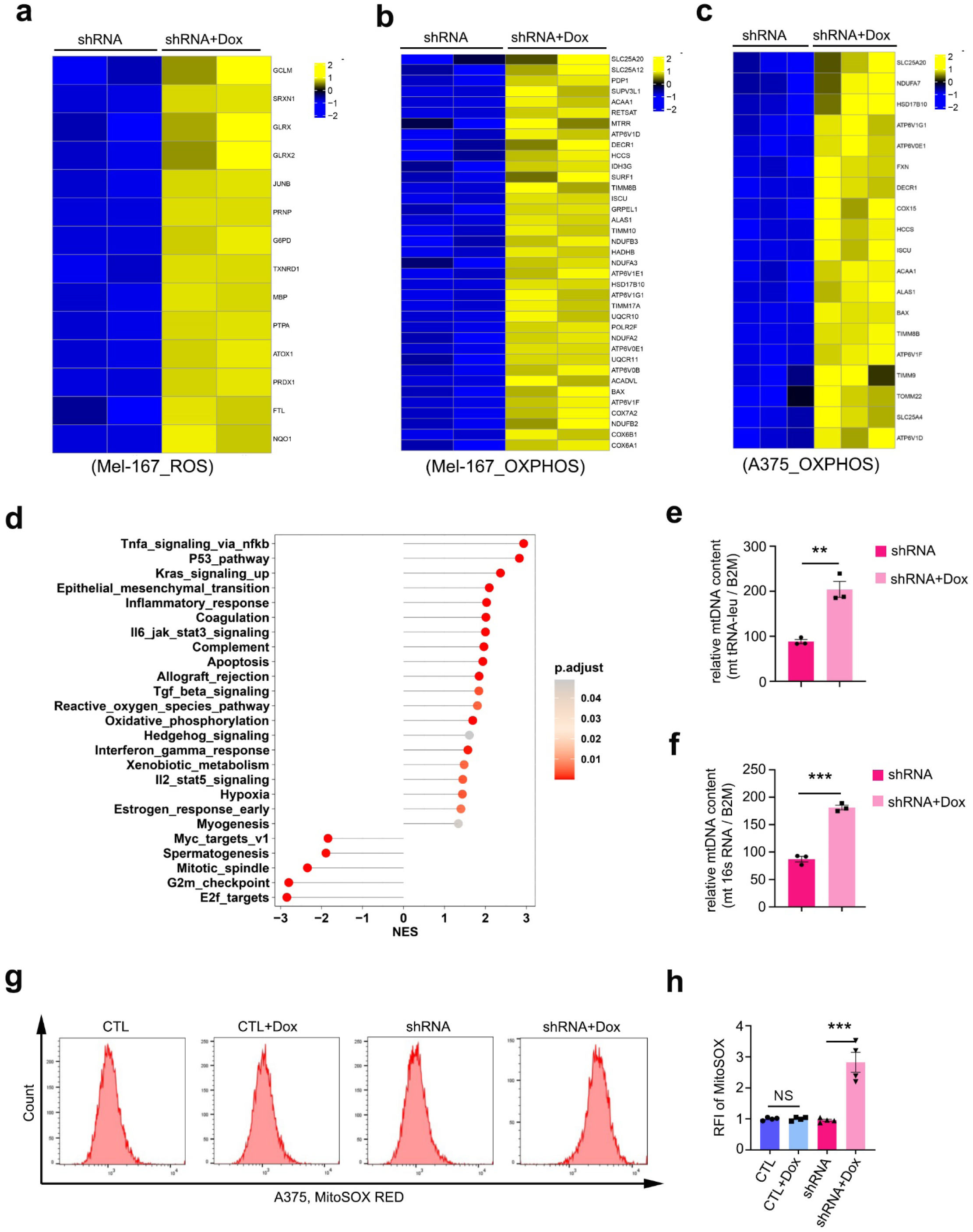
Cortactin depletion results in mtROS elevation. **a**, Heatmap showed the upregulated differential expression gene of ROS pathway of Mel-167 with CTTN depletion. Doxycycline treatment for 4 days. Cutoff, *P* values <0.05, log2FC >0.58. n=2 biological repeats. **b, c**, Heatmap showed the upregulated differential expression gene of OXPHOS of Mel-167 (**b**) and A375 (**c**) with CTTN depletion. Doxycycline treatment for 4 days. Cutoff, *P* values <0.05, log2FC >0.58. n=2 and 3 biological repeats for (**b**) and (**c**), respectively. **d,** The lollipop plot showed GSEA analysis of Hallmark50 gene sets between control and CTTN KD in Mel-167. NES, normalized enrichment score. Doxycycline treatment for 4 days. **e, f**, Relative mitochondrial DNA content determined by qPCR, by normalizing mitochondrial tRNA-leu (**e**) or mitochondrial 16s rRNA DNA (**f**) to nuclear B2M DNA level of A375 cells. n=3 intendent experiments. **g, h**, Mitochondrial superoxide of A375 upon CTTN KD for 3 days, detected by MitoSOX Red staining followed by flow cytometry analysis (**g**). Bar graph (**h**) showed the quantification of relative fluorescence intensity (RFI) of MitoSOX Red. n=3 intendent experiments. Data are represented as mean ± SEM in (**e,f**) and (**h**). The statistical significance was calculated by two-sided Student’s t test in (**e,f**) and (**h**). ** *P*<0.01; *** *P*<0.001; NS, not significant.

**Extended Data Fig. 10.**
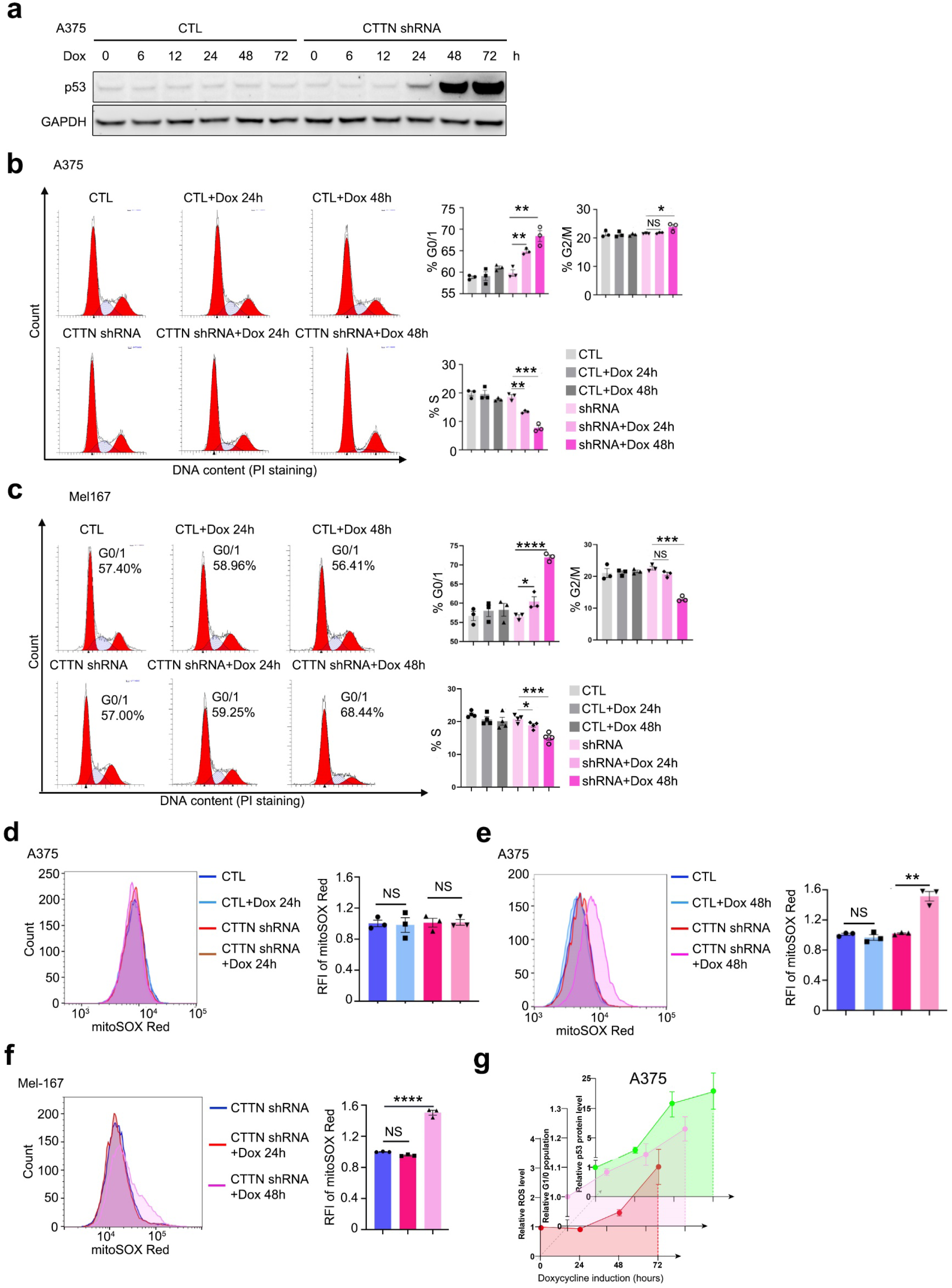
Cortactin depletion induces time-course dependent p53 activation and mtROS elevation. **a**, Immunoblotting showed the p53 protein level changes upon CTTN KD for indicated time in A375. **b, c**, Cell cycle analysis of A375 (**b**) and Mel-167 (**c**) by Propidium Iodide (PI) staining. Cells were treated with doxycycline for indicated time to KD CTTN. Percentage of G0/1 phase, S phase and G2/M phase was quantified. n=3 independent experiments. **d, e**, Mitochondrial superoxide level of A375 cells with or without doxycycline induction for 24h (**d**) or 48h (**e**) determined by MitoSOX red staining followed by flow cytometry. Bar graph showed the quantification of mitochondrial superoxide level. n=3 independent experiments. **f**, Mitochondrial superoxide level of Mel-167 cells with or without doxycycline induction for 24h and 48h determined by MitoSOX red staining followed by flow cytometry. Bar graph showed the quantification of mitochondrial superoxide level. n=3 independent experiments. **g,** Quantification of mitochondrial superoxide (mtROS), % of G0/1 phase and p53 protein level of A375 upon CTTN KD for 24h, 48h and 72h. n=3 independent experiments. Data are represented as mean ± SEM in (**b-g**). The statistical significance was calculated by two-sided Student’s t test in (**b-g**). * *P*<0.05; ** *P*<0.01; *** *P*<0.001; **** *P*<0.0001; NS, not significant.

**Extended Data Fig. 11.**
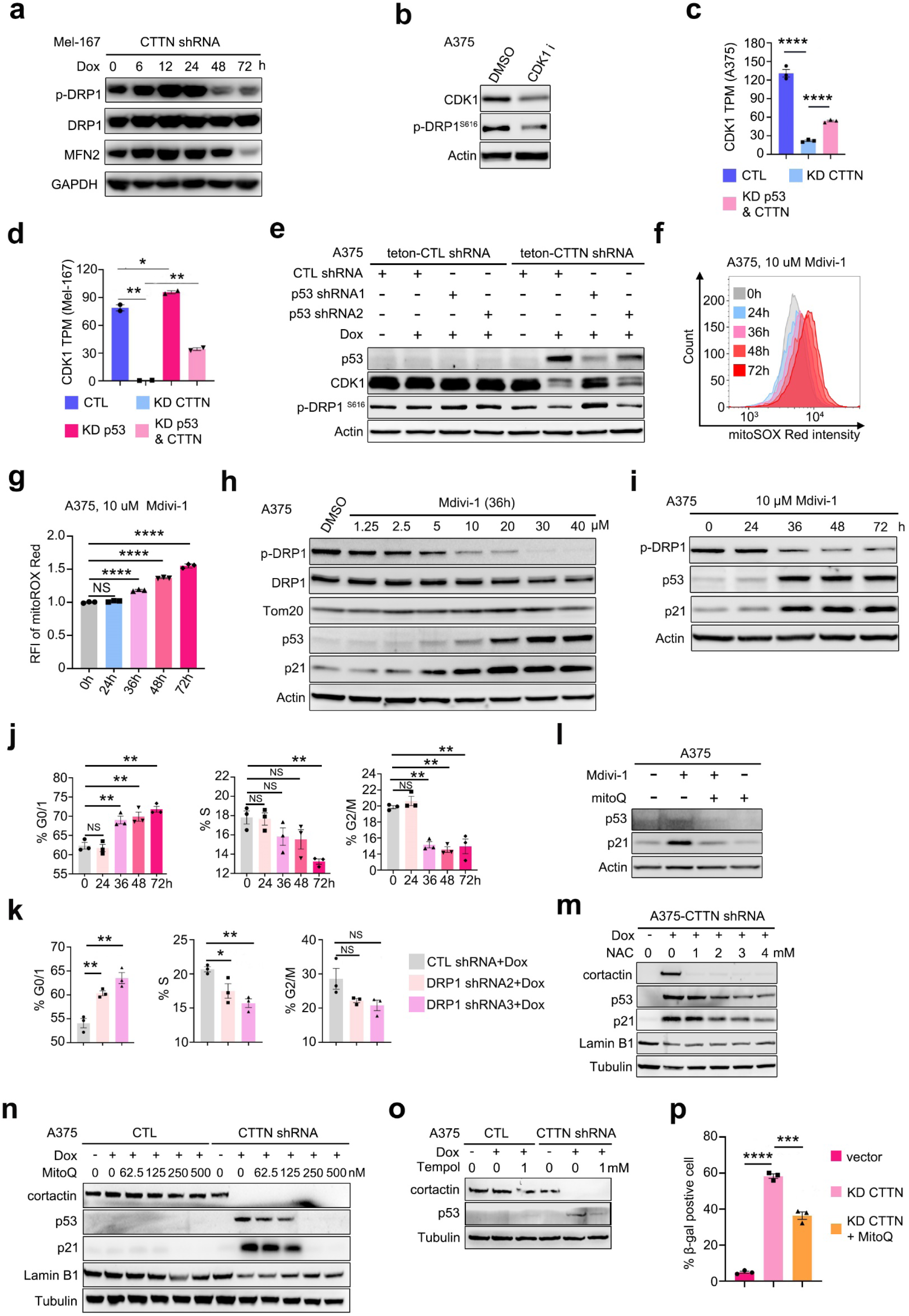
p53 activation and mitochondrial ROS form a signal amplification loop driving stable senescence. **a**, Immunoblotting showed p-DRP1 ser616, DRP1, MFN2 protein level of Mel-167 with or without doxycycline induction to KD CTTN for indicated time. **b**, Protein level of CDK1 and p-DRP1 ser616 determined by immunoblotting. A375 cells were treated with CDK1 inhibitor, 10 μM Cucurbitacin E for 12h. **c,d** , Bar graph displayed the normalized mRNA level (TPM) of CDK1 from RNA-seq data of A375 (**c**) and Mel-167 (**d**) cells. Doxycycline treated for 4 days. n=3 biological repeats for A375 (**c**) and 2 biological repeats for Mel-167 (**d**). **e**, Immunoblotting showed p53, CDK1, p-DRP1 S616 protein level of A375 with or without CTTN or p53 knockdown. CTTN shRNA was cloned into teton-pLKO.1_Puro vector while p53 shRNA was cloned into pLKO.1_Blast vector. **f**, MitoSOX red staining of A375. Cells were treated with 10 μM Mdivi-1 for indicated time. **g**, Quantification of mitochondrial superoxide level determined by mitoSOX red staining of (**f**). n=3 independent experiments. **h**, Immunoblotting determining the protein level of p-DRP1 ser616, DRP1, p53 and p21 of A375 treated with or without DRP1 inhibitor Mdivi-1 for 36h. **i,** Immunoblotting determining the protein level of p-DRP1 ser616, p53 and p21 of A375 treated with or without 10 μM DRP1 inhibitor Mdivi-1 for indicated time. **j**, Quantification of the percentage of G0/1 phase of A375 cells with 10 μM Mdivi-1 treatment for indicated time. **k**, Quantification of the percentage of G0/1 phase of A375 cells with or without DRP1 knockdown. Doxycycline was used to induce the expression of DRP1 shRNA. **l**, Protein level of p53 and p21 of A375 cells treated with or without 10 μM Mdivi-1 and or 250 nM mitoQ treatment for 48h. **m-o**, p53, p21 and or Lamin B1 protein level of A375 pretreated with or without doxycycline to KD CTTN for 1 day then treated with NAC (**m**), MitoQ (**n**) or Tempol (**o**) treatment for 48h. **p,** Percentage of SA-β-gal stained positive A375 cells with or without CTTN depletion and 250 nM MitoQ treatment. n=3 independent experiments. Data are represented as mean ± SEM in (**c,d), (g), (j,k) and (p**). The statistical significance was calculated by two-sided Student’s t test in (**c,d), (g), (j,k) and (p**). * *P*<0.05; ** *P*<0.01; *** *P*<0.001; **** *P*<0.0001; NS, not significant.

## Notes

### Competing Interest Statement

The authors have declared no competing interest.

